# Transformation of coral communities subjected to an unprecedented heatwave is modulated by local disturbance

**DOI:** 10.1101/2022.05.10.491220

**Authors:** Julia K. Baum, Danielle C. Claar, Kristina L. Tietjen, Jennifer M.T. Magel, Dominique G. Maucieri, Kim M. Cobb, Jamie M. McDevitt-Irwin

**Affiliations:** Department of Biology, University of Victoria, PO BOX 1700 Station CSC, Victoria, British Columbia, V8W 2Y2, Canada; Hawaii Institute of Marine Biology, University of Hawaii, Kaneohe, HI, 96744, USA; School of Aquatic and Fisheries Sciences, University of Washington, Seattle, WA, USA; Department of Forest and Conservation Sciences, University of British Columbia, 2424 Main Mall, Vancouver, British Columbia, V6T 1Z4, Canada; School of Earth and Atmospheric Sciences, Georgia Institute of Technology, Atlanta, GA, USA; Hopkins Marine Station, Stanford University, 120 Ocean View Blvd, CA, 93950, USA

**Author notes:** Corresponding Author: Julia K. Baum.

## Abstract

Corals are imminently threatened by climate change-amplified marine heatwaves. Yet how to conserve reef ecosystems faced with this threat remains unclear, since protected reefs often seem equally or more susceptible to thermal stress as unprotected ones. Here, we disentangle this apparent paradox, revealing that the relationship between reef disturbance and heatwave impacts depends upon the focal scale of biological organization. We document a heatwave of unprecedented duration that culminated in an 89% loss of coral cover. At the community level, losses hinged on pre-heatwave community structure, with sites dominated by competitive corals—which were predominantly protected from local disturbance—undergoing the greatest losses. In contrast, at the species level, survivorship of individual coral colonies typically decreased as local disturbance intensified, illustrating that underlying chronic disturbances can impair resilience to thermal stress at this scale. Our study advances understanding of the relationship between climate change and local disturbance, knowledge of which is crucial for coral conservation this century.

## INTRODUCTION

Marine heatwaves threaten the persistence of tropical scleractinian corals (*1–3*), and with them the biologically diverse ecosystems these foundational reef-building species support. Corals are particularly vulnerable to temperature anomalies, with increases of only 1°C capable of disrupting their obligate symbiosis with the photosynthetic dinoflagellate microalgae (family Symbiodiniaceae (*1*)) that normally fuel them, causing the coral animal to expel its symbionts and bleach (*4, 5*). Prolonged bleaching typically leads to coral starvation and mortality (*5*). Although climate change has long been recognized as a serious threat to tropical corals (*6–8*), the recent preponderance of marine heatwaves—persistent anomalously warm ocean temperatures (*9*)—has shifted focus from the threats posed by gradually rising temperatures and ocean acidification (*8*) to these punctuated disturbances (*1, 3*). Already, three global coral bleaching events triggered by El Niño-fueled marine heatwaves (1997–1998; 2010; 2014–2017) have caused devastating coral losses (*10, 11*). Climate change models project that both the intensity and frequency of marine heatwaves will increase in the coming decades (*1, 3*), such that many of the world’s coral reefs are predicted to undergo annual bleaching events by mid-century (*12*). None of these events will, however, occur in isolation. On almost all reefs, climate change is superimposed on a suite of chronic local anthropogenic disturbances (*13*)—ranging from coastal development and associated pollution and reef sedimentation to overexploitation, destructive fishing practices, and disease—that have already significantly altered coral communities through reductions in coral cover and changes to community composition, with largely unknown consequences for species and ecosystem resilience to thermal stress.

Given the intensification of marine heatwaves and the ubiquity of local anthropogenic disturbances on coral reefs, there is an urgent need to understand how these stressors interact (*14*). Yet to date, few coral bleaching studies have explicitly examined multiple stressors (*15*). Chronic local anthropogenic disturbance might mediate coral reef responses to thermal stress, either increasing susceptibility—as documented for massive corals on the Mesoamerican Reef following the 1998 El Niño (*16, 17*)—or conversely enhancing resilience, if disturbances have already eliminated the most vulnerable coral species, leaving behind only the hardiest species (*18*)—as documented on Kenyan reefs (*19*). Alternatively, exposure to chronic local disturbance may have no effect, with thermal stress impacting corals irrespective of underlying protection, as found recently on the Great Barrier Reef (*20*). The degraded state of most modern reefs is widely acknowledged (*8, 13, 21, 22*), and as managers seek to understand how to manage coral reefs under climate change, one might expect that examination of multiple stressors would be common practice in modern coral reef research. However, when we systematically reviewed studies reporting on the effects of recent marine heatwaves (2014–2021)—a period that includes six of the seven hottest years on record—we found that only 10% (n = 20/194) had explicitly tested if local anthropogenic disturbance (or conversely, local protection) influenced heat stress effects on corals (fig. S1, supp. data S1). Approximately half of those that did (n = 9/20) reported a positive effect of protection and increased survival of corals during heat stress events, while 40% (n = 8/20) reported no effect and 15% (n = 3/20) of studies stated that protection reduced coral resilience to heat stress. Conflicting evidence amongst the few bleaching studies that have tested for the effects of local anthropogenic disturbance, and the overall lack of attention to this fundamental aspect of modern coral reefs, impedes understanding of how best to manage these ecosystems in a warming world.

How coral reefs are transformed by climate change this century will depend not only on their exposure to thermal stress and local anthropogenic disturbance but also on the sensitivity and response capacity of individual coral species to these stressors (*23–25*). Coral sensitivity to thermal stress is determined by biological traits, such as tissue thickness (*26*), and physiological tolerance, which is influenced by factors including the type and abundance of the coral colony’s obligate algal endosymbionts (*27–30*). Response capacity, in turn, may reflect species-specific propensity for acclimatization (e.g., the flexibility to switch or shuffle symbionts or to upregulate host thermal stress responses) and adaptation (e.g., selection for traits of either the host or symbionts that confer a fitness advantage under stressful conditions) (*28, 31, 32*). Interspecific differences in sensitivity to thermal stress have long been recognized (*23, 26, 33*), with ‘winners’ generally able to either avoid bleaching during thermal stress or recover from it after warming subsides, and ‘losers’ tending to bleach and die quickly in response to warming (*34*). Since environmental filtering is stronger under stressful conditions (*35–37*), reefs increasingly stressed by marine heatwaves may lose diversity and converge towards simpler assemblages as ‘losers’ are eliminated from the species pool. However, because repeated heatwaves may turn some ‘winners’ into ‘losers’ and vice versa (*38*), questions remain about how corals with different sensitivities will respond to heatwaves of increasing frequency, duration and intensity. Which corals will endure in communities will also depend upon whether species exhibit positive or negative co-tolerance to thermal stress and local anthropogenic disturbance (*35, 39*). Predicting future reef states thus requires not only accounting for underlying anthropogenic disturbances but also understanding interspecific variability in survivorship through heatwaves. To date, however, most heatwave studies have focused on quantifying coral bleaching, a symptom of thermal stress, rather than coral mortality, a fundamental parameter required to quantify the demographic effects of such events. This disconnect reflects the challenge of quantifying coral mortality, which under the strictest standards requires following individual colonies over time, and at minimum requires quantifying coral cover before and after a heatwave, as opposed to bleaching assessments which require only a single site visit. Although bleaching may be an accurate proxy for mortality in short heatwaves, during prolonged events that are becoming the norm, for corals that either bleach quickly and die (and hence are unlikely to be recorded in the bleached state) or those that can persist in a bleached state for prolonged periods, it will not (*40*).

Here, we took advantage of the ecosystem-scale natural experiment created at the epicenter of the 2015–2016 El Niño, the central equatorial Pacific Ocean, where prolonged heat stress blanketed a spatial gradient of chronic local anthropogenic disturbance on the world’s largest atoll, Kiritimati. We quantified thermal stress around the atoll using high-precision *in situ* temperature loggers and satellite data. Our primary objective was to evaluate if protection from local disturbance modulates the impacts of stress on corals. We sought to examine this question both at the community level (i.e., amongst coral species) and at the species level (i.e., for individual coral species). In addition, we evaluated if coral bleaching, the most commonly recorded reef metric during heatwaves, accurately predicts coral mortality. Thus, over the course of nine expeditions (2013–2017) before, during, and after the heatwave, at sites exposed to varying levels of local disturbance (Fig. 1, A and D, fig. S2, S3), we quantified coral community composition and bleaching (n > 250,000 points from 94 photo surveys) and, in one of the largest longitudinal studies of individual corals to date (*41*), tracked the fate of > 850 individual coral colonies.

**Fig. 1.**
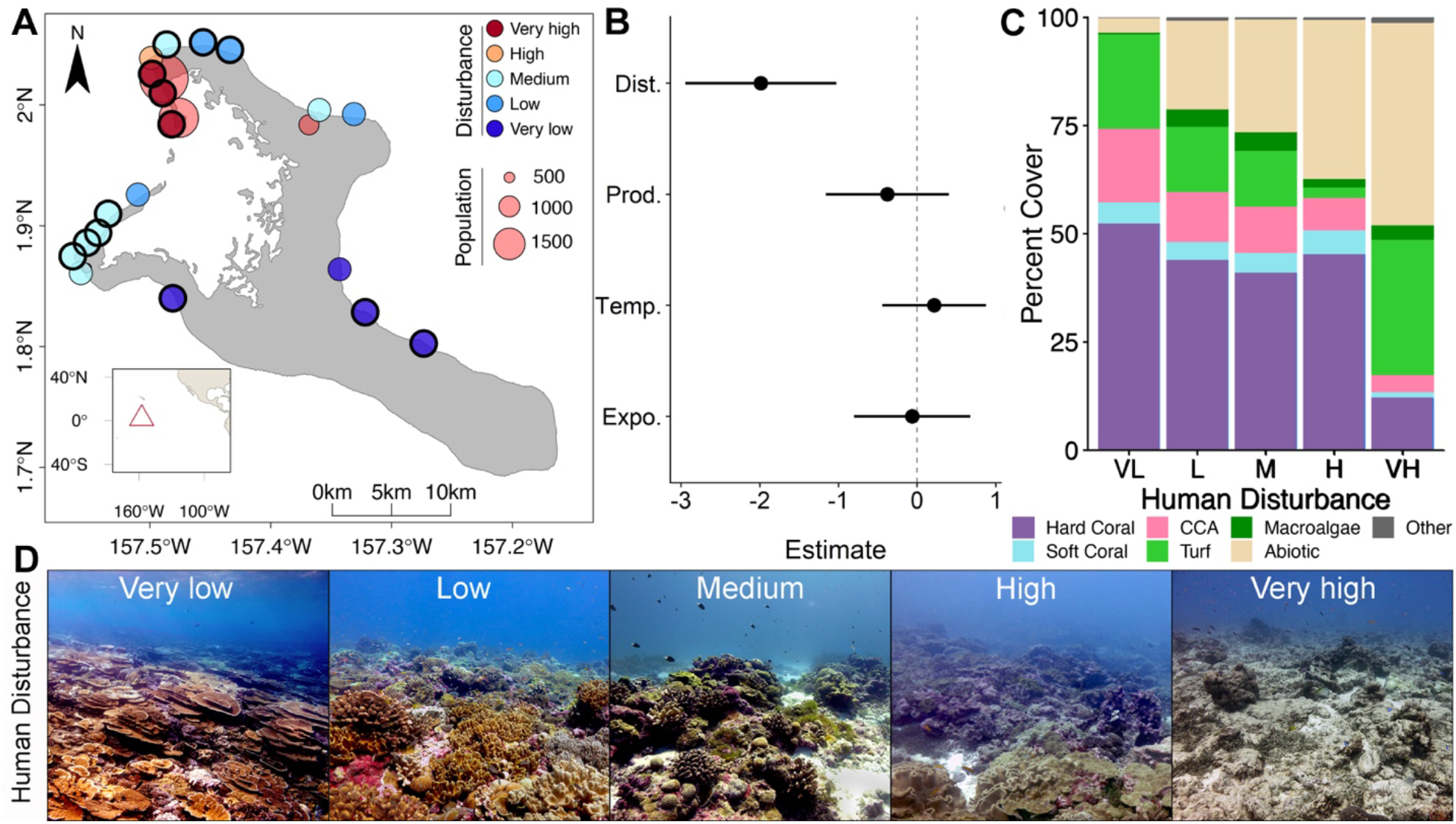
Reef communities across a gradient of chronic human disturbance, prior to thermal stress. **(A)** Reef sites on Kiritimati (central equatorial Pacific Ocean) at which coral community structure and individually tagged coral colonies (sites encircled in black) were tracked over the course of the 2015–2016 El Niño; **(B)** parameter estimates and 95% confidence intervals for factors examined (Dist. = local human disturbance, Prod. = net primary productivity, Temp. = temperature, Expo. = wave exposure) for their influence on hard coral cover prior to thermal stress; **(C)** mean community composition (CCA = crustose coralline algae; turf = turf algae; abiotic = sediment, sand, rubble) of the forereef benthos amongst sites, classified by their exposure to chronic local human disturbance; **(D)** photos of the coral reef communities prior to the El Niño, at sites representing each of the atoll’s levels of local human disturbance.

## RESULTS

### Prolonged heat stress superimposed on reefs spanning a local disturbance gradient

Prior to the El Niño, benthic communities varied dramatically across the atoll’s forereefs, with sites ranging from a high of 62.7% hard coral cover to a low of only 1.6% (Fig. 1). Chronic local human disturbance (fig. S2, S3, table S1; detailed in Supplementary Materials), including dredging and pollution, was the primary determinant of these differences, with coral cover declining significantly as local disturbance increased (*z* = −4.063, *P* < 0.001; Fig. 1B). Abiotic factors, including oceanographic productivity, site exposure (windward versus sheltered), and sea surface temperature, did not significantly influence coral cover amongst sites (Fig. 1B, table S2). Reefs far from villages, with very low exposure to chronic local human disturbance, were amongst the most pristine remaining on the planet before the heatwave, with almost three-quarters of their benthos composed of hard and soft corals and beneficial crustose coralline algae (mean = 52.4 ± 13.2% (SD) hard coral cover) (Fig. 1C). In contrast, reefs exposed to the highest levels of local human disturbance had little hard coral cover (12.2 ± 17.3%), with most of the benthic community composed of turf algae (31.3 ± 18.6%), sediment (28.4 ± 15.5%), sand (11.9 ± 3.2%), and rubble (5.1 ± 1.7%) (Fig. 1C). The effects of different intensities of chronic human disturbance on reef states were strikingly evident visually prior to the El Niño-induced heatwave (Fig. 1D).

As the epicenter of the 2015–2016 El Niño, Kiritimati’s coral reefs experienced a sustained heatwave (Degree Heating Weeks (DHW, °C-weeks) > 0) for approximately one year (Fig. 2). Heat stress started accumulating on 17 April 2015 and exceeded 4 °C-weeks (NOAA’s Coral Reef Watch (CRW) Bleaching Alert 1) from 28 May 2015 until 13 April 2016; DHWs > 0 persisted until 17 July 2016 (Fig. 2A). Accumulated heat stress rapidly exceeded both NOAA’s CRW Bleaching Alert Level 2 threshold (8 °C-weeks) and its 12 °C-weeks threshold, reaching an unprecedented level (> 24.7 °C-weeks; Fig. 2A, table S3) by January 2016. Heat stress was remarkably consistent around the atoll, varying by a maximum of only 4.3% (1.08 °C-weeks) across sites over the course of the event (fig. S4, table S3). Maximum temperature anomalies ranged across sites from 2.83°C to 3.05°C above the reef’s normal maximum monthly mean (MMM) temperature during the event (table S3). The heat stress sustained by Kiritimati’s reefs during this El Niño far exceeds that at any other time point from the recent past for which there are records (fig. S5).

**Fig. 2.**
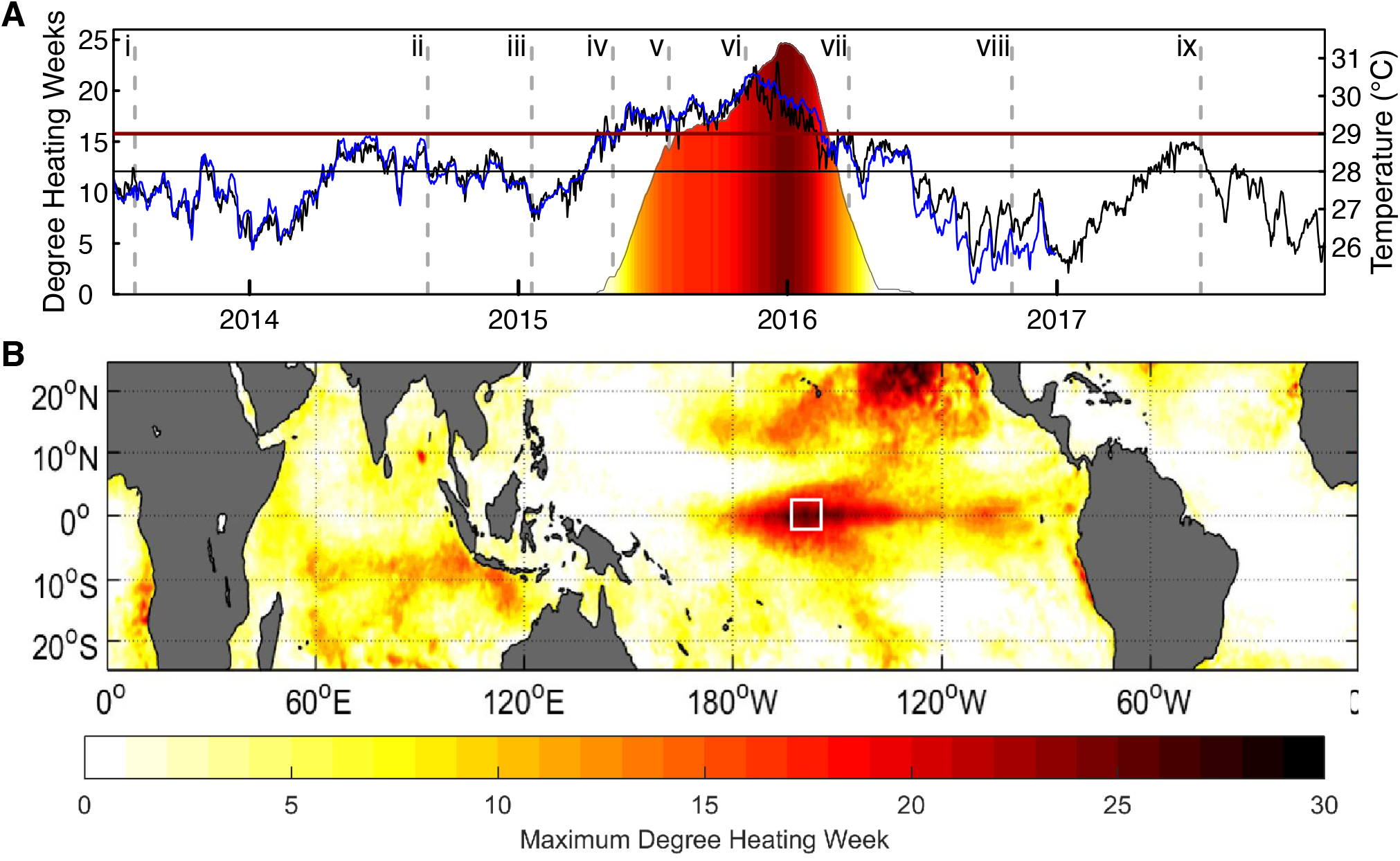
Thermal stress at the epicenter of the 2015–2016 El Niño. **(A)** *In situ* (blue line) and satellite (black line; from NOAA (*42*)) temperature on Kiritimati’s reefs, with maximum monthly mean (MMM) temperature and bleaching threshold for reference (black and red horizontal lines, respectively; from NOAA CRW (Coral Reef Watch) (*43*)); all right axis. Color shows cumulative heat stress on Kiritimati as degree heating weeks (DHW, °C-weeks; left axis), from NOAA CRW (*43*). Dashed vertical lines denote the timing of expeditions prior to (i–iv), during (v–vii), and after (vii–ix) the event. **(B)** Global heat stress on coral reefs during the 2015– 2016 El Niño (May 2015–June 2016) from NOAA CRW, with white box denoting Kiritimati’s location at the epicenter of the heat stress during this event. Color (scale at bottom) indicates maximum thermal stress (°C-weeks).

### Prolonged heatwave impacts primarily reflect differences in coral community composition

The exceptional heat stress unleashed on Kiritimati during the 2015–2016 El Niño caused staggering coral mortality, culminating in an overall estimated loss of 89.3 ± 7.1% of the forereef’s hard coral cover (Fig. 3A, 4). Consistency in thermal stress around the atoll meant that DHW was not a significant factor explaining variability in overall hard coral cover amongst sites (*z* = −0.447, *P* = 0.655; table S4). Nor was bleaching prevalence early in the heatwave (fig. S6) related to the final overall loss of coral cover in the community (*z* = −0.633, *P* = 0.527). Instead, we found that heat stress period (i.e., before vs. after the event; *z* = 18.096, *P* < 0.001) and local disturbance (*z* = −5.146, *P* < 0.001) were both significant predictors of coral cover. The interaction between local disturbance and heat stress period was also significant, indicative of the absolute loss of corals being much greater at minimally disturbed sites than at those exposed to very high disturbance (Fig. 3B), which resulted in the strong inverse relationship between coral cover and local disturbance being completely eroded by the end of the heatwave (*z* = 6.538, *P* < 0.001; slope = −0.73, 95% CI: −1.66–0.21; Fig. 3B, table S4). Relative coral cover losses also tended to be greater with lower local disturbance: on average, minimally disturbed sites underwent an estimated 91.9 ± 1.9% decline in coral cover, ending the heatwave with only 4.7 ± 1.5% coral cover, while sites exposed to very high levels of local disturbance—which already had depressed levels of coral cover—declined by a further 64.6 ± 15.1%, ending the heatwave with only 3.89 ± 4.6% coral cover (Fig. 3B, 4). However, this difference in relative coral cover losses was not quite statistically significant (*t* = −3.1124, *P* = 0.09), likely due to variable losses at the high local disturbance sites. Reef-building corals were replaced primarily by turf algae, which rapidly overgrew the dead coral, and at some sites also by macroalgae (Fig. 4, fig. S7).

**Fig. 3.**
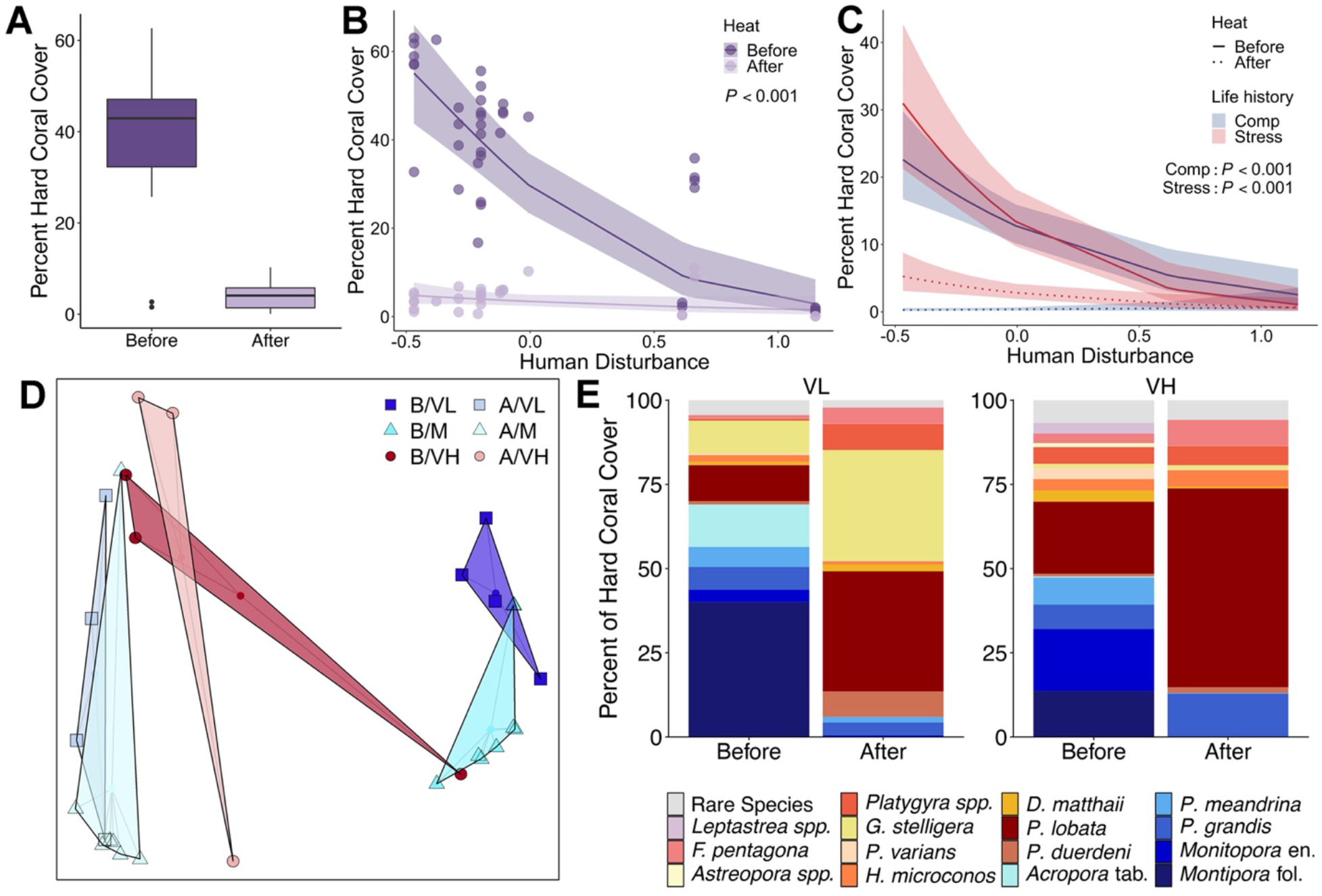
Impact of a prolonged heatwave on coral community composition. **(A)** Overall change in hard coral cover across all sites from before to after the 2015–2016 El Niño on Kiritimati (sites as in Fig. 1A); model predictions of the effect of local human disturbance on percent coral cover before versus after the heatwave for **(B)** the overall coral community and **(C)** stress-tolerant (red) and competitive (blue) corals; **(D)** PCoA plots of coral assemblage structure before (B) and after (A) the event at very low (VL), medium (M), and very high (VH) levels of local human disturbance; and **(E)** comparison of average coral community composition across sites exposed to very low disturbance versus those exposed to very high levels of disturbance, before and after the heatwave (stress-tolerant species in shades of red – yellow, competitive species in blues).

**Fig. 4.**
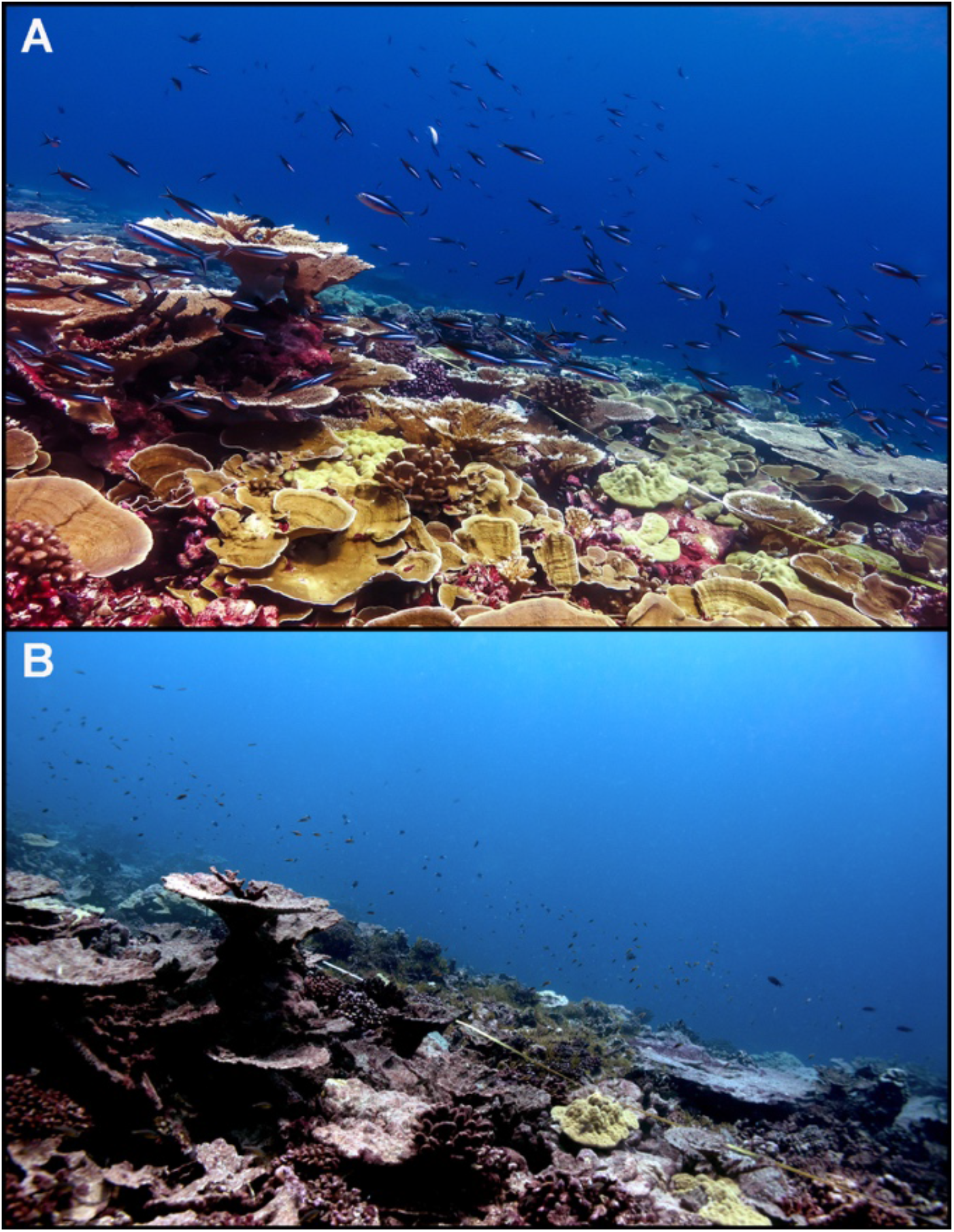
Transformation of a minimally disturbed coral reef by a heatwave of unprecedented duration. **(A)** Before (July 2015) and **(B)** after (July 2017) the 2015–2016 El Niño-induced mass coral mortality event at one site (VL1) with very low exposure to chronic local stressors on Kiritimati. Virtually all coral in (B) is dead and overgrown by turf algae. Photo credits: (A) Kieran Cox, University of Victoria and (B) Kristina Tietjen, University of Victoria.

We hypothesized that differences in coral cover loss across the local disturbance gradient reflected distinct coral communities, comprised of species with variable thermal stress tolerances, found at different disturbance levels. Examining these coral communities revealed that not only did community composition vary with disturbance prior to the heatwave (PERMANOVA, *pseudo-F* = 4.4, *P* < 0.001; Fig. 3D, dark shaded polygons), but that it also changed significantly as a result of it (PERMANOVA, *pseudo-F* = 15.3, *P* = 0.002), with sites exposed to very low or medium local disturbance experiencing greater turnover than those exposed to very high disturbance (PERMANOVA, *pseudo-F* = 3.9, *P* = 0.022; Fig. 3D). Underlying this change was the loss of corals with a competitive life history strategy, namely all large tabulate and corymbose *Acropora* at very low local disturbance sites (100% loss) and all foliose *Montipora* at very low and medium local disturbance sites (100% loss; Fig. 3E, 4). Models testing the relationship between pre-heatwave community composition and coral cover showed that sites dominated by ‘competitive’ corals were more strongly impacted by the heatwave than those dominated by ‘stress-tolerant’ corals (*z* = −3.169, *P* = 0.002; fig. S8). Moreover, separate models of competitive and stress-tolerant coral cover revealed that while the cover of both life histories decreased significantly due to the heatwave (competitive: *z* = −13.252, *P* < 0.001; stress-tolerant: *z* = −12.293, *P* < 0.001; Fig. 3C, table S4), the degree of change was substantially greater for competitive corals (96.8 ± 4.8%) than for stress-tolerant ones (78.7 ± 10.5%; *t* = −6.8807, *P* < 0.001). We also found that, unlike for overall hard coral cover, the magnitude of thermal stress at each site (°C-weeks) was statistically significantly related to the cover of both competitive and stress-tolerant corals; however, its effect size was much smaller than that of local disturbance or heat stress period (Table S4).

### Chronic local disturbance can exacerbate heatwave impacts on individual coral species

In contrast to the observed changes at the community level, at the species level we found a clear signal of the negative influence of chronic local disturbance on coral survival through prolonged heat stress (Fig. 5; fig. S9). Examining a representative sample of the same seven coral species across sites showed a significant negative relationship between local disturbance and coral survival (*z* = −3.79, *P* < 0.001; Fig. 5A, table S5). This signal became clearer when distinguishing between life history strategies, with survival of the stress-tolerant coral species strongly negatively related to local disturbance (Fig. 5, B, D and F) while that of the competitive coral species showed no relationship with disturbance, namely because these colonies were so sensitive to the heat stress that few of them survived (Fig. 5, C, E, and F; 2.0% survival for *Pocillopora grandis;* 2.97% for *Montipora aequituberculata*). Individually, each stress-tolerant coral species exhibited an inverse relationship between survival and local disturbance, with the massive corals *Platygyra ryukyuensis* and *Porites lobata* exhibiting the steepest relationships (an estimated 95% and 80% survival at very low sites, and 4% and 29% survival at very high sites for the two species, respectively) and *Hydnophora microconos* the shallowest (58% survival at very low sites to 28% survival at very high sites) (Fig. 5F, Fig. S9, S10, Table S5). Survivorship of the two competitive species each showed weakly positive, but non-significant, relationships with local disturbance, with mortality at all sites exceeding 97.5% (Fig. 5F, Fig. S9, S10). Bleaching prevalence early in the heatwave was not related to the ultimate survival of any coral species (*P* > 0.69) (Figs. S11, 12).

**Fig. 5.**
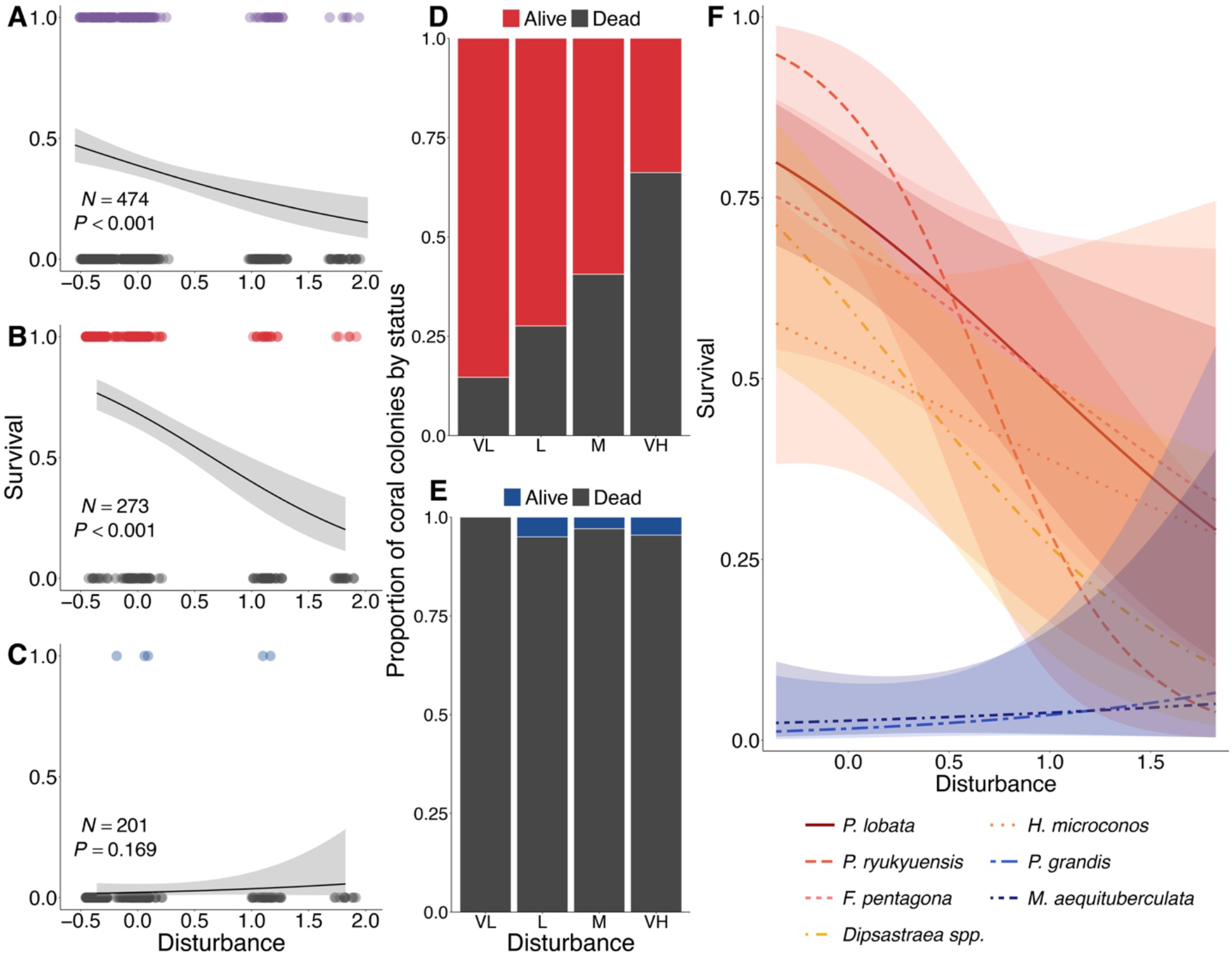
Survival of n = 475 individually tracked colonies throughout the heatwave. Logistic regressions of coral survival versus disturbance for: **(A)** all coral colonies, **(B)** stress-tolerant (red) species, **(C)** competitive (blue) species, and **(F)** all coral colonies, with species modelled as a fixed effect, in a two-way interaction with local disturbance; bar plots of coral colony survival by local disturbance level for **(D)** stress-tolerant and **(E)** competitive coral species.

### Emergent winners and losers

Given strong interspecific differences in survival, certain coral species emerged from this unprecedented heatwave as winners, while others were clear losers (Fig. 6). All of the competitive coral species were losers, having each lost over 97.5% of their cover (Fig. 6A). In contrast, the winners were all stress-tolerant coral species that underwent smaller losses in cover (65.4% to 87.2% for the five biggest ‘winners’) and thus increased considerably in their relative proportion of coral cover (Fig. 6). Notably, the massive coral *Porites* spp., which was already the most common coral prior to the heatwave, underwent more than a two-fold relative increase, such that over half of the atoll’s remaining coral community is now comprised of this one slow-growing, stress-tolerant species (Fig. 6B). Although two other corals—*Platygyra* spp. and *Pavona duerdeni—*had even greater proportional gains, because they were initially relatively rare, they still comprised only a small proportion of the coral community at the end of the heatwave (10.6% and 3.1%, respectively; Fig. 6B). Finally, substantially lower rates of colony mortality than coral cover loss for the five individually tracked stress-tolerant species reflects the fact that while many colonies sustained partial mortality during the heatwave, a portion of the colony was still alive at the end of it (Fig. 6A).

**Fig. 6.**
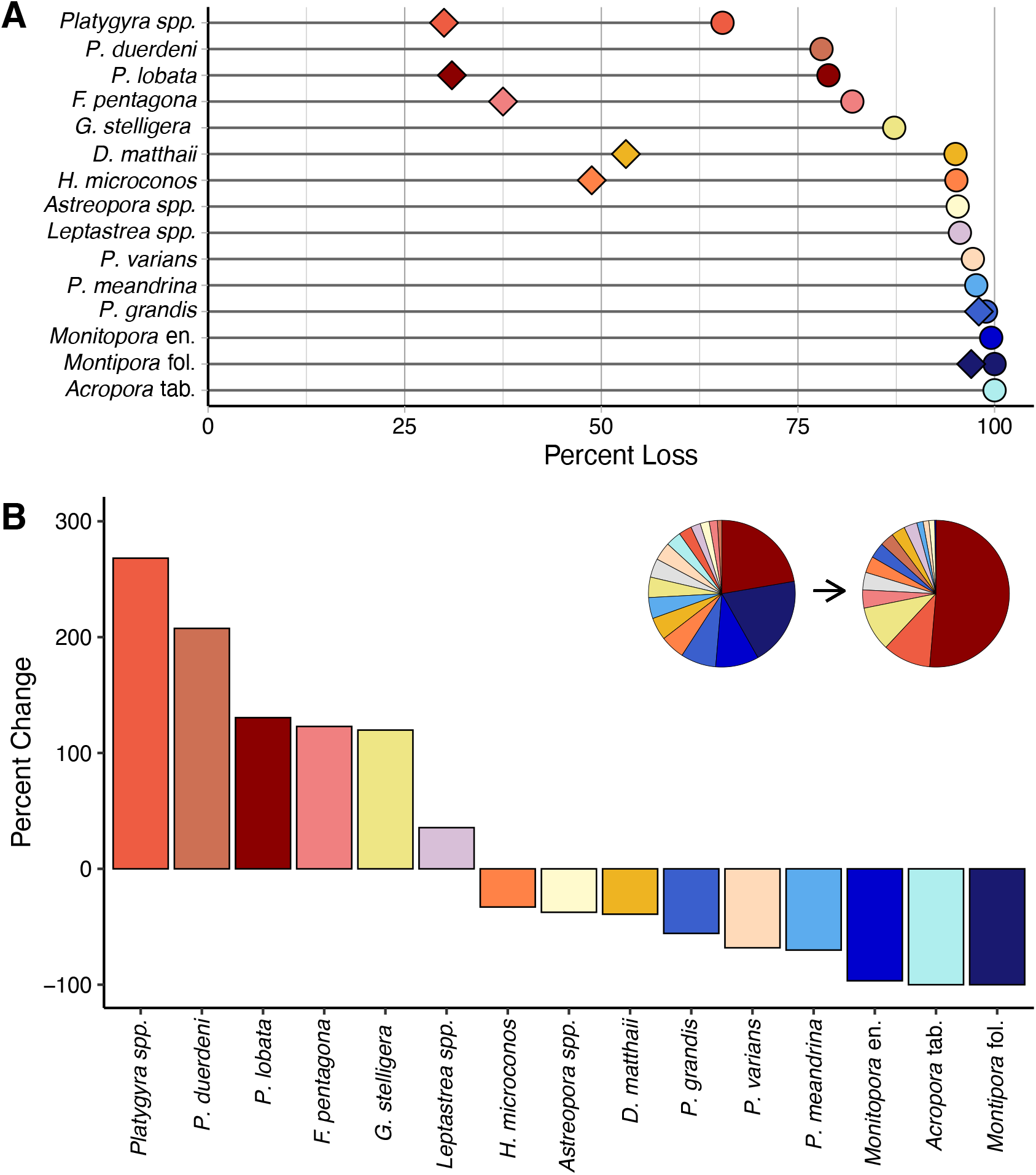
Interspecific variation in prolonged heat stress impacts on corals. **(A)** Overall change in percent cover of individual coral taxa (circles) for the 15 taxa comprising the greatest proportion of benthic cover prior to the heatwave, ordered from least to greatest cover loss; overall percent colony mortality for the seven individually tracked coral species is overlaid (diamonds). **(B)** Percent change in proportion of overall coral cover, from before to after the heatwave, ordered left to right from species that underwent the greatest proportional gains (‘winners’) to the greatest proportional loss (‘losers’). Inset pie charts show species composition of the overall coral assemblage before (left) and after (right) the heatwave (grey = rare species). See table S6 for taxonomic and other details.

## DISCUSSION

Despite the urgent need to leverage all available solutions for enhancing coral reef resilience to thermal stress in the face of escalating climate change, it has been unclear if protecting reef’s from underlying local anthropogenic disturbance helps or hinders in this regard. Our study sheds new light on this debate. Tracking whole coral communities and individual species exposed to consistent thermal stress, but different levels of underlying local anthropogenic disturbance, clarified that the relationship between local anthropogenic disturbance and climate resilience varies qualitatively across biological scales. At the community level, we found that reefs exposed to high levels of chronic local anthropogenic disturbance fared better through a prolonged heatwave than those shielded from local disturbance, an outcome driven by differences in the coral community composition amongst sites. In contrast, when comparing survival rates within individual coral species, the predominant relationship for those species that were not eradicated by the heatwave was one of declining survivorship as local disturbance increased. These findings have implications for managing and restoring coral reefs as climate change-driven marine heatwaves continue to intensify.

### Winners and losers in prolonged heatwaves

Overall, we documented mass coral mortality (89% hard coral cover loss) arising from a heatwave that persisted for a remarkable ten months unabated. At its peak, heat stress accumulated to 25 °C-weeks (degree heating weeks), a level that had not previously been anticipated to occur on any reef until mid-century (*26*). Its occurrence 35 years earlier than predicted underscores how rapidly climate change is advancing (*1, 2, 9*). Although corals typically exhibit high interspecific variability in their sensitivity to thermal stress (*33*), a heatwave this extreme might have been expected to overwhelm the tolerance of even the most resilient species, resulting in high mortality rates across the board. Instead, we found that interspecific coral cover losses still varied widely, from the complete loss of tabular *Acropora* and foliose *Montipora* to a loss of only 65% in the stress-tolerant mounding coral *Platygyra* spp. Mortality rates for the individual colonies we tracked were also highly variable across species, but with lower mortality rates for the stress-tolerant species because many of the colonies of these species had surviving corallites, thus potentially paving the way for their recovery. Only on nearby tiny Jarvis Island was thermal stress more extreme (maximum 31.58 °C-weeks) during this heatwave (*44, 45*). There, reefs underwent an estimated >98% decline in hard coral cover, with severe losses of *Montipora* spp. (100%), *Pocillopora* spp. (>90%), and *Pavona* spp. (~85%) recorded in the surveyed areas (*45, 46*). Together, these results illustrate that extremely prolonged heatwaves will still have ‘winners’ and ‘losers’, such that these events will cause not only dramatic coral losses but also substantial changes in community composition.

### Influence of local anthropogenic disturbance on coral survival

In species-specific models of the influence of exposure to chronic local anthropogenic disturbance on coral resilience to thermal stress, we detected an inverse relationship for those species with ‘stress-tolerant’ life histories. Survivorship of all stress-tolerant species was at least twice as high at sites sheltered from local stressors compared to those with the highest exposure, and over 10 times as high for that with the strongest inverse relationship, *Platygyra ryukyuensis*. Increased coral sensitivity to thermal stress is, we expect, most likely attributable to the diminished water quality at the most highly disturbed sites. On Kiritimati, raw sewage and pollution inputs have resulted in increased turbidity and sedimentation (Fig. S3) and greater concentrations of bacteria, virus-like particles, and potential pathogens in the water column at these sites (*47, 48*). Previous studies have shown that low water quality can change the coral microbiome (*49–51*), and microbiome analyses of subsets of our tracked corals prior to the heatwave showed increased bacterial diversity at highly disturbed sites (*52*). Such changes can have knock-on effects for coral physiology and survivorship even in the absence of warming (*53*), which may then be exacerbated during heatwave events. Poor water quality, as with other environmental stressors, can also lead to changes in coral–algal symbioses (*54–56*), which may then influence coral resilience to subsequent thermal stress. Indeed, an analysis of our tracked *Platygyra ryukyuensis* colonies revealed that distinct *Symbiodiniaceae* genera across the disturbance gradient were linked to coral survival during heat stress (*57*). Although the mechanism underlying the relationship between coral survival and local disturbance remains unclear for the other stress-tolerant corals we tracked—and may differ across species—we suggest that it likely results from distinct coral microbiomes, and their associated physiological traits, found across the disturbance gradient. We attribute the failure to detect an effect of local disturbance on coral resilience to thermal stress in our two competitive coral species to their extremely high mortality. A recent study of a less severe heatwave, on Moorea, French Polynesia, showed that bleaching severity was significantly increased by local nitrogen pollution in the competitive coral genera *Acropora* and *Montipora* (*58*). Overall, these results suggest that impairment of coral resilience to thermal stress by local stressors may be a general phenomenon, and at least deserves increased research and management attention.

Strikingly, however, these species-level results stand in contrast to our own community-level results, and previous studies, which have suggested that local disturbance enhances coral reef resilience to thermal stress. Over a decade ago, Côte and Darling (*59*) argued that this would be the case for coral reefs exposed to, and altered by, local stressors, because the most sensitive coral species would already have been eliminated, leaving behind a more stress-tolerant community. That is, if organisms exhibit co-tolerance to stressors (or, conversely, co-sensitivity) such that they respond similarly to them, then the combined effects of the stressors may be antagonistic, resulting in a response that is less than the sum of their individual effects (*14, 39, 60*). Although such a response is not a given, with multiple stressors also sometimes eliciting synergistic or additive responses, it appears not uncommon on reefs exposed to local stressors and global climate change. Darling et al. (*19*) found that while the stress-tolerant corals that dominated fished reefs in Kenya prior to the 1998 El Niño were barely impacted by the bleaching event, reefs in no-take reserves had more diverse coral assemblages, including many corals with competitive life-history traits that exhibited co-sensitivity to fishing and bleaching, and incurred heavy losses. More recently, Cannon et al. (*61*) showed that central Pacific reefs in the Gilbert Islands that were exposed to higher levels of chronic local pressure were dominated by a coral species tolerant of nutrient loading and turbidity and were subsequently less impacted during a bleaching event than nearby reefs with fewer local pressures. At Kiritimati’s highest disturbance sites, we found that of the competitive coral species, *Acropora* were completely absent, and while some encrusting *Montipora* persisted, only a few colonies of the foliose form (common in less disturbed sites) were recorded. Thus, the ‘positive’ effect of local disturbance reflects different community compositions and the variable thermal sensitivities of the coral species that dominate disturbed reef communities, rather than there being a mechanism by which local disturbance itself enhances coral resilience to thermal stress.

More difficult to reconcile with either our species- or community-level findings are studies reporting that coral responses occur irrespective of local protection, influenced only by the reefs exposure to thermal stress (*18*). In surveys of the Great Barrier Reef Marine Park during the 2016 marine heatwave, for example, Hughes et al. (*20*) documented severe bleaching on reefs in each of the park’s types of management zones, and concluded that local management of water quality and fishing pressure had little to no influence on coral resistance to extreme heat. Similarly, a study in one of Indonesia’s oldest marine parks during the same heatwave found that management zone made no difference to coral losses (*62*). More recently, Baumann et al.’s (*63*) global meta-analysis tested the relationship between human influence and coral resilience and concluded that reefs isolated from human pressures are not more resilient to climate change, noting that even the world’s most remote reefs bear the impacts of intense marine heatwaves. We concur that at broad spatial scales, exposure to thermal stress will be highly variable across the considered reefs, and this may well be the primary determinant of reef impacts; remote reefs are not immune to high thermal stress exposure levels. Such an emphasis on current and future thermal stress exposures has proven useful when considering future thermal refugia for coral reefs, as in the ‘50 Reefs’ conservation prioritization (*64, 65*). At finer spatial scales, however, where thermal stress exposure is the same (or very similar) across reefs, a corollary of the conclusion that coral responses are (or appear to be) the same irrespective of protection is that coral sensitivities and response capacities to thermal stress must be the same across the protection levels. We can envision only a few means by which these conditions could be met: *1*) corals are exposed to similar conditions inside and outside the protected area, such that the communities do not differ. This is likely to be the case in some areas where MPAs either have inadequate enforcement or have not been established long enough for coral recovery to have occurred within the MPA; *2*) exposures to thermal stress across protection levels were not actually equal; *3*) failure to detect different impacts due to insufficient power, or measuring bleaching at only a single time point such that the full ecological impacts of the event were not quantified; or *4*) the coral communities differed because of the stressor, but the coral species in the different areas had equal sensitivities and response capacities to heat stress. Scenarios one to three do not imply that local disturbance has no effect on coral resilience to thermal stress, and the high interspecific variability in thermal tolerance makes the latter scenario unlikely.

Considering our results together across scales suggests that, although local anthropogenic disturbance can result in the loss of sensitive coral species such that the remaining community is more tolerant to subsequent thermal stress, when comparing ‘apples with apples’—that is, the same species across different levels of local anthropogenic disturbance—there is clear evidence that local disturbance can impair survival. Thus, while there is compelling recent evidence that coral reef recovery following bleaching events may not be aided by reef protection (*18*), our study suggests that previous conflicting results pertaining to coral community resilience to thermal stress can be resolved through consideration of biological scale.

### Coral bleaching does not foretell demographic impacts of prolonged heatwaves

Our repeated reef surveys during an extended bleaching event also provide an empirical test of the relationship between coral bleaching and mortality. Given the many challenges associated with conducting *in-situ* assessments of coral bleaching events—including the need to marshal resources quickly when heatwaves arise, the limited reef area that can be assessed by divers, and the complexities of accessing remote reefs—rapid reef surveys at a single time point during a heatwave are often used to assess ecological impacts. But whereas bleaching incidence may be a reliable indicator of diminished coral fitness (given that it can lead to decreased coral growth and reproduction), the capacity of corals to recover from bleaching means that it may not accurately foretell coral mortality, and hence the demographic impacts of heatwaves. Indeed, we found no relationship between bleaching prevalence and subsequent mortality levels in any of our tagged coral species. Instead, we found that the species with the highest bleaching incidence early in the event (*Platygyra ryukyuensis* and *Favites pentagona*) had amongst the lowest mortality, while a species with very low bleaching incidence (*Pocillopora grandis*) suffered near complete mortality (Figs. 6A, S11, S12); results were similar for our benthic community data. Mismatches between bleaching and mortality could arise if certain coral species can resist the onset of bleaching more than others, but then only persist in a bleached state for a short period (*40, 66*). Such mismatches will be more likely in the prolonged heatwaves that are predicted to become more common under climate change (*40, 67*), thus highlighting the need for increased sampling during these events to accurately gauge demographic impacts. As the capacity to use satellite-derived data to accurately monitor coral bleaching increases, these sources could help to overcome this challenge.

### Coral reef recovery from prolonged heatwaves unlikely under climate change

We posit that coral reef recovery from prolonged heatwaves is increasingly unlikely, because of long ecosystem recovery times and the diminishing interval between successive heatwaves under climate change (*68, 69*). On Kiritimati, our sampling up until three years after the end of the heatwave (2019, prior to the onset of COVID-19) revealed new juvenile corals and regrowth of colonies that had experienced partial mortality, which together resulted in some increase in overall coral cover but still left the ecosystem a long way from full recovery. Long-term studies of coral reefs from the Indian and Pacific Oceans following the major 1998 El Niño found that recovery of hard coral cover typically took more than a decade and involved substantial turnover of community composition, with ‘recovered’ reefs tending to have lower coral diversity and be dominated by fast-growing corals (*70–73*). Recovered reefs in Moorea, for example, are now dominated by ‘fields’ of weedy *Pocillopora*, while recovering reefs in the Seychelles became dominated by fast-growing, branching *Acropora* corals (*74*). Reef recovery following mass bleaching events is also not guaranteed. Following the 1998 El Niño, over 40% of surveyed reefs in the Seychelles underwent regime shifts to fleshy macroalgae (*73*). Those that were on a recovery trajectory, which had high coral cover prior to the 1998 El Niño, still had not fully recovered by 2014, and although full recovery was projected to be complete within 17 to 29 years (*74*), progress was nullified by Seychelles’ 2016 bleaching event (*75*). Such outcomes are increasingly likely with climate change (*68*). Thus, as with many reefs, full recovery of Kiritimati’s reefs now seems unlikely.

Persistence of coral reefs throughout the 21^st^ century will be dictated almost entirely by the extent to which greenhouse gas emissions are reduced (*67*). Our study shows that prolonged heatwaves under climate change will not only significantly reduce coral cover but also transform coral community composition. Diminishing intervals between recurrent heatwaves will leave most reefs with insufficient time to recover (*68*). Emissions reductions that only limit warming to 2°C are projected to result in the loss of virtually all coral reefs (99%), whereas if warming is limited to 1.5°C, losses could be limited to between 70% and 90% (*67, 76*). Under such dire conditions, strategies additional to GHG emissions reductions that can reliably enhance coral resistance to, or recovery from, marine heatwaves should be broadly deployed. Yet, the efficacy and scalability of the potential options remains uncertain. Our study provides evidence that coral species’ resilience to thermal stress is enhanced as local anthropogenic stressors are reduced. These findings imply that alleviating local stressors—such as by improving water quality, which is likely one of the most tractable options for reef managers—could not only benefit natural coral reefs but also aid coral restoration efforts, improving the odds of success for the individual coral species that are out-planted on reefs. With much still to learn about the interactions between multiple stressors on coral reefs, we encourage researchers to explicitly incorporate local stressors into future studies of marine heatwave impacts on coral reefs. In addition to urgent reductions in greenhouse gas emissions, evidence-based local management actions that are both scalable and durable are urgently needed as a means of increasing the odds of persistence for these imperiled ecosystems under climate change.

## MATERIALS AND METHODS

### Literature survey

We conducted a systematic review of the primary literature to quantify the extent to which field studies assessing the impacts of recent marine heatwaves (i.e., 2014 to 2021) on corals quantified underlying local anthropogenic disturbance at their study site and tested for an effect of them on coral outcomes through the heat stress event. On 9 September 2021, we conducted a search for papers published between 2015 and 2021 using all databases on the Web of Science, with the following search terms: ((“coral*”) AND (“mortal*” OR “bleach*” OR “cover*” OR “health*”) AND (“El Niño” OR “El Nino” OR “ENSO” OR “heat*” OR “thermal stress” OR (“temperature” AND “anomal*”) OR “bleaching event”)). We evaluated each of the n = 721 papers returned from this search, reviewing the titles, abstracts, and method sections, to first determine if the paper examined corals during a heatwave between 2014 and the present day; we excluded papers describing lab-based studies or heatwaves prior to 2014. Additionally, we added n = 10 papers that were not returned from this search but were known to quantify the effects of a heatwave between 2014 and the present day. The remaining n = 184 papers that met our criteria were classified based upon if the study included an anthropogenic disturbance (searching for “anthropogenic”, “human”, “disturbance”, “stressor”, “cumulative effects” or “protection”) and if the study analyzed or made conclusions about the effect of anthropogenic disturbances. We also noted the type of disturbance (e.g., fishing, pollution), if the study included sites without disturbance as a control, the coral sampling method (e.g., randomized quadrats, etc.), as well as the frequency of sampling before, during, and after the heatwave event.

### Study site

Situated in the central equatorial Pacific Ocean at the center of the Niño 3.4 region (a designation used to quantify El Niño presence and strength) (*77*), Kiritimati (Christmas Island) is the world’s largest atoll by landmass (388 km^2^, 150 km in perimeter). Coral reefs are exposed to vastly different levels of chronic local human disturbance depending on their location around the atoll (Fig. 1A). Human impacts—including pollution from sewage outflow and an oil company, major infrastructure (i.e., a pier), and fishing pressure on the forereef—are densely concentrated on the northwest coast, where the two main villages are located and the majority of the population resides (Fig. 1A, table S1). In contrast, reefs on the atoll’s north, east, and south coasts are minimally impacted (Figs. 1, S2; detailed in Supplementary Materials) (*78, 79*).

We quantified the intensity of chronic local human disturbance at each forereef site (described below in Field Methods), as in (*57, 80*), using two spatial data sources: 1) the number of people residing within 2 km of each site, as a proxy for localized impacts, based upon the Government of Kiribati’s 2015 population census data for each village on Kiritimati (*81*); 2) subsistence fishing pressure, quantified through detailed semi-structured interviews conducted with heads of household in each of the atoll’s villages in 2013 (*82*) and represented using a kernel density function as a measure of its intensity at each site. We combined these data with equal weight to create a quantitative metric of chronic local human disturbance at each site (table S1). This metric correlated strongly with sedimentation, turbidity, and bacterial loads, three other indicators of disturbance (see Supplementary Methods, Fig. S2). For visualization purposes, and to contrast reefs exposed to local disturbance extremes, we also classified each site as one of five distinct disturbance levels (very low, low, medium, high, and very high) based upon clear breakpoints in our continuous disturbance metric (Fig. 1A, table S1) (*57, 80*). These terms should be regarded as being relative to other levels of disturbance around the atoll, rather than absolute levels of human disturbance.

In addition to local human disturbance, we also quantified site-specific oceanographic parameters to assess their influence on benthic community composition around the atoll (detailed in Supplementary Methods). We extracted remotely-sensed data for maximum net primary productivity and wave energy from the open-source data product Marine Socio-Environmental Covariates (MSEC (*83*)) and defined site exposure (i.e., windward versus leeward) based on the dominant wind direction (southeasterly (*84*)).

### Field methods

To examine how heat stress interacts with chronic local human disturbance, we conducted benthic surveys at nineteen forereef sites on the atoll during nine expeditions between 2013 and 2017: four prior to the onset of thermal stress (July 2013, August 2014, January and May 2015), three during the El Niño-induced heat stress (July and November 2015, March/April 2016), and two after the event (November 2016, July 2017). The surveyed reefs at each site are all at 10–12 m depth on sloping, fringing reefs, with no back reef or significant reef crest formations, and adjacent sites are all more than 1 km apart (with one exception) (*85*). On average, we surveyed 10.4 ± 4.9 sites per expedition, for a total of 94 surveys (table S7); logistical and weather constraints associated with working in such a remote location prevented surveying all sites in each time point (table S7).

To survey sites, we photographed the benthos underneath a 1 m^2^ gridded quadrat (mean = 28.1 ± 4.1 quadrats per site) at randomly selected positions adjacent to a sixty meter transect that had been placed along the 10–12 m isobath (n = 2,637 photos total; table S7). Photographs were taken with a Canon Powershot digital camera (G15 and G16 models with an Ikelite housing and wide-angle lens dome) that was white-balanced at depth on each dive. We analyzed all benthic photos using CoralNet, an open-source online software for benthic image analysis (*86*), by projecting 100 random points onto each image and manually identifying the substrate beneath each point (n = 259,359 total) from our custom label set (n = 103 identification tags), which consisted of coral (table S6) and non-coral animals, algae, bacteria, and abiotic substrates, such as sand and sediment. Recognizing that some corals cannot be definitively identified to the species level by morphology alone, we have identified some corals to the genus level only. For each coral taxon, we confirmed that the morphotype was consistent across all sites. We also included separate labels for bleaching and non-bleaching coral tissue (e.g., ‘bleaching *Porites*’, ‘ *Porites*’), thus allowing us to determine the proportion of points per site in each expedition that were bleaching for each coral taxon. We quantified each site’s benthic community composition in each surveyed time point by dividing the total number of points for each substrate type by the total number of annotated points from all quadrats (detailed in Supplementary Methods).

Additionally, we tagged and photographed 834 individual coral colonies from seven species at thirteen of our nineteen monitoring sites (Fig. 1A) and tracked their fate over the course of the El Niño event. We selected three common corals as our focal species (*Porites lobata, Pocillopora grandis, Montipora aequituberculata*) because they were found at all sites and include different life history strategies, with the first classified as being a ‘stress-tolerant’ coral, a group that is defined by slow growing, massive species and the capacity to tolerate chronically stressful and variable environments, and the latter two considered to be ‘competitive’ corals, a group typified by large, branching and plating species with fast growth and the capacity to dominate communities (*19, 87–89*) (table S6). We aimed to tag twelve colonies of each of these three species per site. We also tagged up to six colonies per site of each of four less common species (*Favites pentagona, Dipsastraea* spp. (primarily *D. matthaii*)*, Platygyra ryukyuensis, Hydnophora microconos*), each of which has a stress-tolerant life history strategy (*88*) (table S6). Tagged coral colonies were located along the same transects as the benthic photoquadrats. Colonies were first tagged and photographed during the August 2014 expedition, prior to the onset of heat stress. For each coral, we photographed the entire colony parallel to the colony surface with a ruler next to it for scale; macro shots were taken of the colony surface to aid in identification where necessary. In each subsequent expedition (except November 2015), we re-photographed each colony that could be relocated and also tagged and photographed additional colonies.

We assigned each coral colony a bleaching status for each time point in which it was photographed using the following visual criteria: 1) no bleaching or paling; 2) some light bleaching but less than 5 cm across the largest patch and less than 50% of colony pale; 3) bleaching in patches >5 cm or more than 50% of colony pale; 4) severe or complete bleaching (>80%) or the entire colony pale. For binomial treatments of bleaching, we considered categories 1 and 2 to be “healthy” and categories 3 and 4 to be “bleached”. Thus, colonies were assigned to “bleached” if they had at least one patch of their surface that was bleached and greater than 5 cm across or if more than 50% of the surface of the coral was faded.

In total, we were able to determine the survivorship status of 474 of the tagged colonies (average of n = 36.5 colonies per site; range = 9 to 56; Table S7); the remaining colonies could not be relocated after the heatwave. Corals were recorded as surviving the heatwave if they were found alive at any time point following the event, and as not surviving it if they were found dead upon first inspection post-heatwave. This occurred at the end of the heatwave (March/April 2016) for most colonies, and in the two subsequent expeditions for corals located at sites that we had either been unable to fully sample (i.e., one dive instead of two to three to search for all corals) or sample at all (due to unfavorable weather conditions or logistical constraints) in March/April 2016.

### Temperature and thermal stress

We quantified temperature on Kiritimati during the 2015–2016 El Niño event using both remotely sensed data extracted from NOAA’s Coral Temp product (*42*), as well as high precision *in situ* temperature loggers (Sea-Bird Scientific SBE 56; ± 0.001°C precision). *In situ* loggers were deployed at sites around the atoll (minimum of one logger deployed per disturbance level; n = 17 sites, including n = 12 of the sites surveyed in this manuscript) between 2011 and 2016, all at ~10 m depth (range 8–12 m) on the forereef (fig. S1). For both data sources, we quantified temperature for all available sites around the atoll as in Claar et al. (*90*) and averaged across sites to produce a measure of island-wide temperature.

To assess the potential influence of baseline temperatures on coral communities around the atoll prior to the 2015–2016 El Niño, we also extracted the maximum monthly mean temperature (*91*) for each site from NOAA Coral Reef Watch’s (CRW) monthly mean sea surface temperature climatology, which are produced at a 5-km spatial resolution (ftp://ftp.star.nesdis.noaa.gov/pub/sod/mecb/crw/data/5km/v3.1/climatology/nc/).

We quantified thermal stress on Kiritimati during the 2015–2016 El Niño as degree heating weeks (DHW; °C-weeks), the metric most commonly employed to assess coral bleaching risk. Corals are sensitive to temperatures more than 1°C above their long-term maximum monthly mean (MMM) sea surface temperature (SST), known as the bleaching threshold. DHW is a measure of accumulated thermal stress, which is defined as the rolling sum of temperatures above the bleaching threshold during the preceding twelve weeks (*92, 93*). Significant coral bleaching is expected to occur once cumulative thermal stress has exceeded 4 °C-weeks (NOAA CRW Bleaching Alert Level 1), with widespread bleaching and some mortality typically expected at > 8 °C-weeks (NOAA CRW Bleaching Alert Level 2) (*43*).

DHW values for Kiritimati were extracted from the U.S. NOAA CRW’s 5-km DHW product (NOAA CRW Daily Global 5-km Satellite Coral Bleaching Heat Stress Degree Heating Week Version 3.1) (*93*) for January 2011–December 2016 (*90*), for each of the nineteen study sites, and used to calculate an island-wide mean (Table S3). Comparisons of these satellite-derived thermal stress values to *in situ* estimates in a previous study (*90*) yielded consistent results. Herein, we present the satellite-derived DHW data and analyses employing these data for comparability with other coral bleaching studies. Additionally, to compare the thermal stress experienced on Kiritimati during this heatwave to earlier events, we extracted DHW values (as above) from 1985 to 2018 (Fig. S4).

### Statistical analyses

Analyses were conducted in R 4.0.4. interfaced with Rstudio 1.4.1106.

We fit a series of generalized linear mixed-effects models (GLMMs) with the benthic community composition data, in which the overall proportion of hard coral cover (i.e., the response variable) was modelled with a beta error distribution and a logit link, using the *glmmTMB* package (*94*). First, to examine influences on coral cover prior to the 2015–2016 El Niño, we modelled overall hard coral cover as a function of chronic local disturbance (continuous) and three environmental variables: maximum net primary productivity (NPP; mg C m^-2^ day^-1^), sea surface temperature, and site exposure (windward vs. sheltered). Second, to assess the influence of prolonged heat stress on coral communities, and if chronic local human disturbance modulates heat stress impacts, we modelled overall hard coral cover as a function of heatwave period (before vs. after), maximum heat stress experienced during the El Niño (i.e., maximum site-level DHW), chronic local disturbance, and a two-way interaction between heatwave period and disturbance. Third, we examined if the impact of the heatwave on coral cover was modulated by the pre-El Niño coral community composition. To do so, we defined a ‘dominant coral life history’ covariate for each site by classifying each coral species according to its life history strategy (following (*88*), table S6), then classifying each site as ‘stress-tolerant dominated’ (if > 60% of the corals at that site had this life history strategy), ‘competitive dominated’ (if > 60% of the corals at that site had this life history strategy), or ‘mixed’ (if there was no dominant life history type). We included this covariate as a fixed effect in a model of overall hard coral cover, with a two-way interaction between it and heatwave period (before vs. after); maximum site-level DHW was also included as a fixed effect. Fourth, to directly quantify the impacts of heat stress on corals with distinct life history types, we fit separate models for the cover of stress-tolerant corals and the cover of competitive corals. In these models, coral cover was modelled as a function of heatwave period, chronic local disturbance (including the two-way interaction with heatwave period), and maximum site-level DHW. We fit two different versions of these life history models: one where the cover of each life history type was calculated as the proportion of overall benthic cover and one where it was calculated as the proportion of total hard coral cover. In all models, continuous explanatory variables were standardized using the ‘rescale’ function in the *arm* package, and site was included as a random effect to account for the non-independence of data collected at the same site over time. Expedition was also included as a random effect in all models (with site and expedition modelled as crossed random effects), except for the competitive coral cover model, as its inclusion in this model led to convergence issues. To test the sensitivity of the competitive model to this change, we ran all the other models without expedition as well and found that this did not result in any significant changes to the model results. See Supplementary Methods for additional details.

We also employed a multivariate approach to examine differences in hard coral community composition across the disturbance gradient, both before and after the heatwave, by conducting multivariate ordinations and statistical analyses using the *vegan* package (*95*). A site-by-species matrix was created for the entire hard coral community using measures of percent cover. We performed multivariate ordinations (principal coordinates analysis; PCoA) using the ‘betadisper’ function to visualize differences in the coral communities amongst the three most disparate (very low, medium, very high) levels of local human disturbance and across heat stress periods. We then tested for significant differences in coral community structure using permutational multivariate analysis of variance tests (PERMANOVA; ‘adonis’ function) with 999 permutations and Bray–Curtis distances. Heat stress, human disturbance, and their interaction were included as fixed effects, while site was incorporated as a blocking factor using the strata term in ‘adonis’.

We used our longitudinal tagged coral dataset to directly examine the impact of prolonged heat stress on the survival of individual coral colonies. In all cases, coral survival was modelled using generalized linear models (GLMs), with a binomial distribution and logit link, in the *stats* package. We modelled the survival of stress-tolerant and competitive corals separately, with coral species, chronic local disturbance, maximum site-level DHW, maximum net primary productivity, and site exposure (windward vs. sheltered) included as fixed effects in each model, plus a two-way interaction between coral species and local disturbance. We present the results of these models, and visualizations of these models without species included (to show the relationship between coral survival of each life history strategy and disturbance; Fig. 5, B, and C), as well as an overall model (all species, with a species-by-disturbance interaction, to show the relationship between survival of all seven coral species with disturbance; Fig. 5F). To visualize the overall pattern, we also modelled and displayed all corals together without life history or species in the model (Fig. 5A), but with the other covariates. All continuous explanatory variables were standardized using the ‘rescale’ function in the *arm* package. For each of these model subsets (e.g. overall model, stress-tolerant model, competitive model, etc.), we ran models with all possible combinations of variables. We then used AIC to determine the top model for each model subset. In all cases, the model with disturbance (and the two-way life history or species interactions, where included) but without any of the environmental variables had the lowest AIC. Additionally, in all cases, for the models within 4 ΔAIC of the model with the lowest AIC, disturbance (and the two-way life history or species interactions, where included) was significant, but other variables were not. We initially also included ‘site’ as a random effect, but in all cases this worsened the model fit.

Finally, to assess if bleaching is an accurate metric of heatwave outcomes for corals, we a) used the benthic photoquadrat data to test if the extent of bleaching at a site early in the heatwave (July 2015) was a predictor of the final overall coral cover loss at each site, and b) used the tagged coral colony data to test if corals that exhibited bleaching early in the heatwave had lower survival through the event. First, using the photoquadrat data, we calculated the proportion of hard corals that were bleached in July 2015 at each site (n = 13 sites, as not all sites were surveyed in July 2015; fig. S7) as well as the loss of coral cover. The proportion of coral cover lost at each site was calculated by averaging coral cover values across field seasons within each heatwave period, then using the following formula:

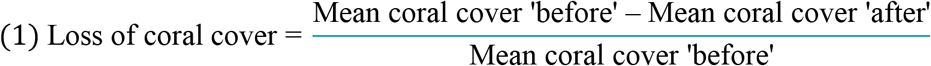

We modelled the loss of coral cover as a function of three continuous variables—proportion of bleached corals, local disturbance, and maximum heat stress (°C-weeks)—using GLMMs with a beta error distribution and logit link. Separate models were fit for the overall hard coral community and for both competitive and stress-tolerant corals. Next, using the tagged coral colony data, we modelled coral survival using GLMs with a binomial distribution and a logit link. Bleaching status, coral species, disturbance, and exposure were included as fixed effects, along with a two-way interaction between coral species and bleaching status. We first treated bleaching as a binary state (as detailed above and reported in the results) and tested the sensitivity of our results to this assumption by also modelling the four different bleaching categories (see Supplementary Materials). We conducted all of these models both for the overall tagged coral dataset, and then for the stress-tolerant and competitive coral species separately. Bleaching results did not differ across any of these different model forms.

## Acknowledgements

We gratefully acknowledge our collaborators and friends on Kiritimati who facilitated this research, R. Bebe, P. Tofinga, T. Kirata, T. Alefaio, V. Hnanguie, F. Tiata, J. Teem, and L. Teem; L. Szostek, M. Watson, S. McNally, K. Cox, T. Stovel, J. Mortimer, J. Burns, K. Bruce for help collecting the field data; N. O’Brien, L. Szostek, R. Hansen, E. Giannantonio and A. Kozachuk for assistance with benthic image processing.

## Author Contributions

Conceptualization: JKB
Data collection: JKB, DCC, KLT, JMI, KMC, DGM
Data processing: DGM
Data analysis: KLT, JMTM, DCC, JKB
Writing—original draft: JKB
Writing—review & editing: JKB, DCC, KLT, JMTM, DGM, KMC, JMI

## Data and materials availability

Data reported in this paper are provided on Zenodo; code to reproduce the analyses and figures is provided on GitHub [https://github.com/baumlab with repository made public upon after acceptance].

