## Supplementary Materials for "Transformation of coral communities subjected to an unprecedented heatwave is modulated by local disturbance"

**This PDF file includes:**

Supplementary Text
Figs. S1 to S12
Tables S1 to S8

**Other Supplementary Materials for this manuscript include the following:**

Data S1

#### Supplementary Methods

Study Site

Kiritimati, an atoll within the Republic of Kiribati, supports approximately 6,500 people (*81*) the vast majority of whom are highly dependent on reef resources for subsistence and income due to the atoll's geographic isolation and limited alternate livelihoods (*13*, *82*). Reef fish in particular are a vital resource, with over 95% of households on Kiritimati actively engaged in fishing activities (*82*). Since 2009, we have monitored forty coral reef sites around the atoll: thirty-seven of the sites were initially established in 2007 by Walsh (*85*) and three new sites were added in 2009. Included herein are the nineteen sites at which we surveyed benthic community composition at least once in the two years before the 2015–2016 El Niño (July 2013–May 2015) and at least once in the year after the event.

*Chronic Local Human Disturbance*

Kiritimati's spatial gradient of chronic local human disturbance arose because of the concentration of villages and infrastructure on the northwest coast (Fig. S2, Table S1). In addition to the villages, there has been extensive dredging for a port near one of the sites on this coast (VH1). Kiritimati also does not have a sewage treatment plant (*47*), and run-off is known to lower water quality on coral reefs (*78*, *79*). Reefs elsewhere on the atoll are subject to minimal local human disturbance. In particular, coral reefs in the Bay of Wrecks and on the eastern side of Vaskess Bay (Fig. S2) experience virtually no direct local human impacts, as there are no villages or infrastructure whatsoever in these areas.

Previous studies have noted Kiritimati's gradient in local human disturbance and the degraded state of Kiritimati's reefs near the atoll's villages. When surveyed in 2007, Walsh (*85*) found that both top predator and carnivorous fish biomass were significantly lower at sites with elevated fish catches (i.e., primarily those categorized herein as 'very high' or 'high' disturbance) compared to those at sites with lower catches (i.e., 'medium', 'low', and 'very low' disturbance sites). Focusing on the microbial and benthic component of the reef ecosystem, Dinsdale et al.'s (*47*) study from across the northern Line Islands, which sampled only reefs across Kiritimati's lagoon face, reported that Kiritimati's reefs had ten times as many microbial cells and virus-like particles in the water column as on uninhabited Kingman Reef. Kiritimati's microbes were reportedly dominated by heterotrophs, including a high proportion of potential pathogens, and the benthic community was said to have the highest prevalence of coral disease of the four surveyed northern Line Islands (*47*). Of the sites sampled on Kiritimati, those in the very high disturbance region (i.e., north of the lagoon) had higher microbial counts and higher counts of culturable *Vibrio* spp. than the site sampled on the south lagoon face (close to our sites that are categorized as medium disturbance) farther from the villages (*47*). More recently, McDevitt-Irwin et al. (*48*) showed that sites on Kiritimati exposed to very high disturbance (VH1, VH2) had significantly higher bacterial counts in the water column (near the coral substrate) compared to sites exposed to very low disturbance (VL1, VL2).

We tested whether our quantitative metric of local human disturbance was correlated with three other indicators of disturbance (sedimentation, turbidity, and microbial load) for our nineteen surveyed sites (Fig. S3). As a proxy for sedimentation, we used our benthic photoquadrat data to calculate an estimate of the percent of the substratum covered by sediment. We calculated this 'percent sediment' metric both including and excluding sand. As a proxy for turbidity, a single experienced scientific diver (K. Tietjen) estimated visibility at each dive site. On expeditions where a site was sampled on more than one day (up to n = 3), these estimates were averaged. We then averaged this visibility across all expeditions for which we had data for a given site. As expected, sites nearest to villages had lower visibility (mean = 14.5 m) than those with the lowest disturbance (mean = 32.3 m; Fig. S2). We tested for relationships between human disturbance and a) percent sediment cover (without sand), b) percent sediment cover (with sand), and c) visibility using linear models. Finally, we re-evaluated the data from McDevitt-Irwin et al. (*48*) on the concentration of bacteria in the water column at four sites on Kiritimati (two very high and two very low human disturbance). These data were collected by taking water samples (1–2 mL), preserving them in formaldehyde and then filtering the samples and counting DAPI-stained bacteria under high magnification. The mean concentrations of microbes at each site (n = 4 samples each) were then compared using a one-way ANOVA and a Tukey post-hoc test (Fig. S2d).

*Oceanographic Factors*

Beyond anthropogenic impacts, natural oceanographic factors including sea surface temperature, oceanographic productivity, and wave energy can influence coral reef ecosystem structure and diversity (*96–99*). To assess the extent to which such features might explain differences in benthic community composition around Kiritimati atoll, we quantified multiple oceanographic and abiotic variables at our nineteen sites (Table S2; water temperature is described in the section below in the context of heat stress). At each site, we quantified *in situ* salinity, dissolved oxygen (DO) saturation, and pH using a YSI Pro Plus handheld multiparameter meter that was calibrated daily, and collected water samples to quantify nutrients (i.e., phosphate, silicate, nitrate, nitrite) (Table S2). We supplemented these *in situ* measurements with remotely-sensed data, using the Marine Socio-Environmental Covariates (MSEC) open source data product (https://shiny.sesync.org/apps/msec/) (*83*) to obtain estimates of oceanographic productivity and wave energy (Table S2), as follows: *1)* maximum net primary productivity (NPP; mg C $m^{-2}$ $day^{-1}$) values in MSEC were calculated over a 2.5 arcmin grid based on data from NOAA CoastWatch, which models NPP using satellite-derived measures of photosynthetically available radiation (PAR), sea surface temperature (SST), and chlorophyll-*a* concentrations; *2)* mean wave energy (kW $m^{-1}$) in MSEC is computed from the WAVEWATCH III hindcast dataset. We excluded the data for sites in which an estimate was made from wind and fetch values rather than the WAVEWATCH III data, detailed in (*83*).

With the exception of primary productivity, which is known to vary across Kiritimati's reefs due to island-wake upwelling that occurs along the atoll's western side (*85*), there was little variation in these oceanographic and water chemistry characteristics amongst sites around the atoll (Table S2). We note, however, that although long-term mean wave energy values at our sites were quite similar, ranging only from ~25 to 27 kW $m^{-1}$, reliable WAVEWATCH III data were not available for several sheltered sites along the lagoon face; thus, these data may not capture the true variability in wave energy across the atoll. We therefore also defined a site exposure variable based upon the predominant wind direction that included all sites (*84*) and employed this variable in our statistical models.

#### Data Processing

##### *Benthic Community Data*

For each photo of the benthic community (n = 2,649; Table S4), we first cropped the image around the quadrat and checked the white balance, adjusting those photos that still had color casts due to technical issues underwater. In CoralNet (*86*), we then manually annotated the substrate beneath each of the 100 random points and identified corals to either genus or species, based on the functional relevance of each coral to the project and our ability to distinguish species from photographs. Points that could not be annotated (e.g., the taxonomic identity could not be resolved because the point fell on a dark shadowed area) were excluded from the dataset for downstream analysis. If a photo had more than 10 points that could not be annotated, it was excluded from the dataset entirely; this resulted in 12 photos (0.42% of photos) being removed from the data set, for a final total sample size of 2,637 (Table S4).

#### Statistical Analyses

##### *Coral Cover Models*

The coral cover models presented in the main text were fit using all available data points, such that some sites had more than one data point per heatwave period because they were sampled in multiple expeditions (Table S4). To test the sensitivity of our results to this imbalance in the data, we ran all the models a second time using a reduced dataset in which values were averaged across expeditions to produce one data point per site for each heatwave period. Models were fit using the same parameters as the main models, excluding expedition (which could no longer be included as a random effect). Using this reduced dataset had minimal effect on the model results for all the models presented (see Table S5).

#### Supplementary Figures

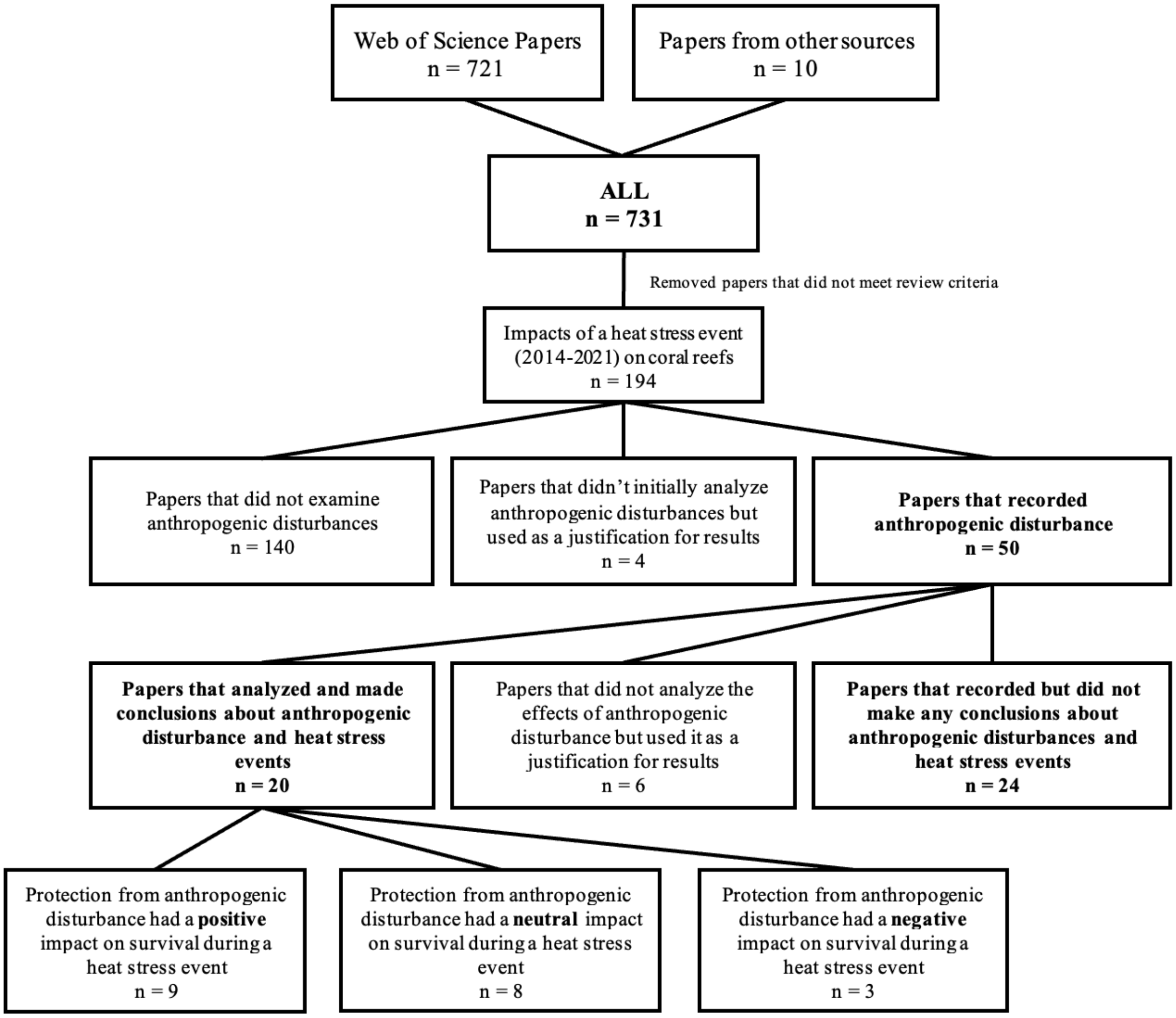


**Fig. S1. PRISMA diagram showing results of the systematic review of the primary literature** to quantify the extent to which field studies that had assessed the impacts of recent marine heatwaves (2014 to 2021) on corals quantified underlying anthropogenic stressors at their study site and tested for an effect of them on coral outcomes through the heat stress event.

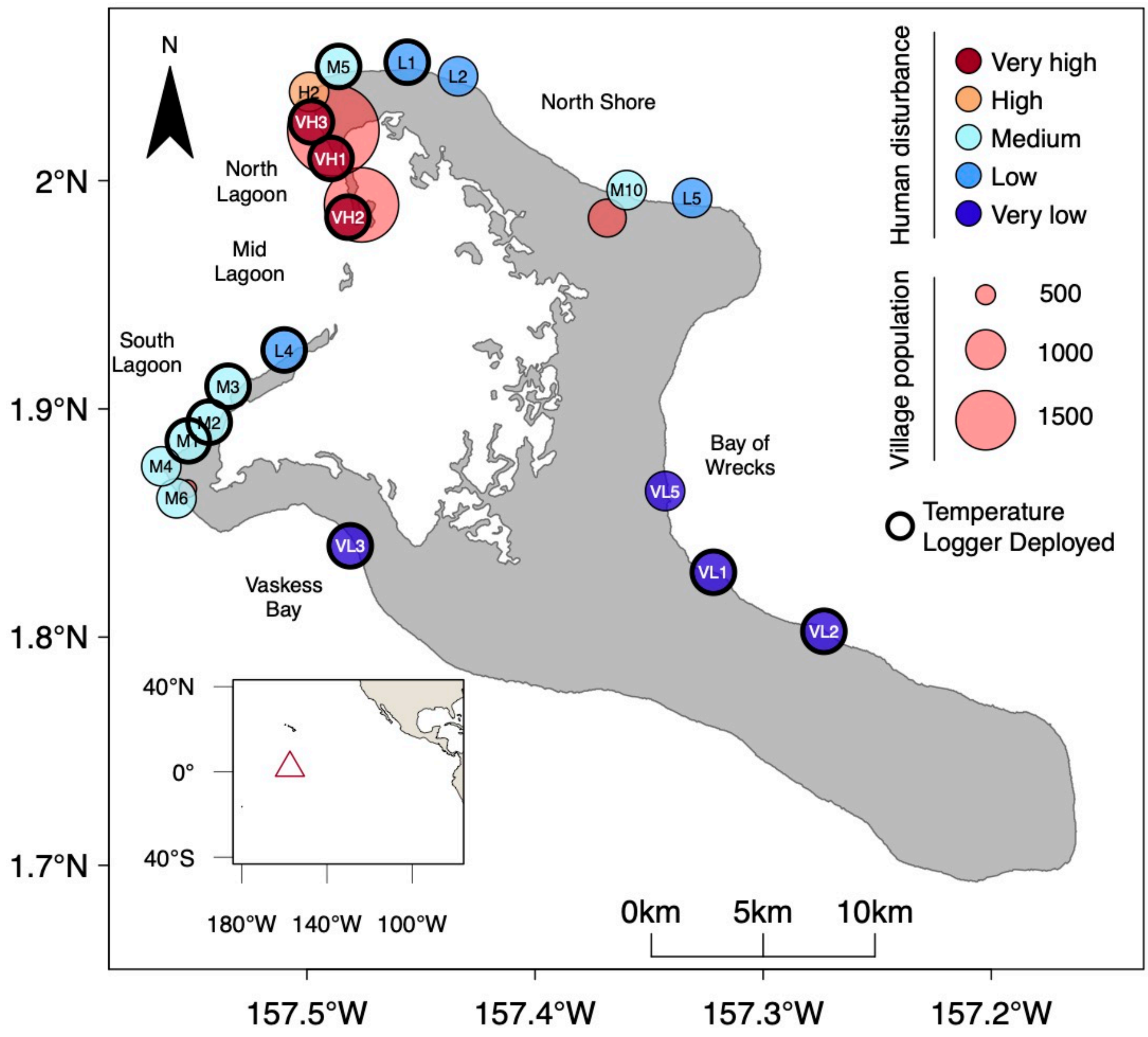


**Fig. S2. Kiritimati (Christmas Island), showing nineteen forereef sites** (and their site names, as in Tables S1–S3) at which benthic community composition was quantified between 2013 and 2017. Sites are categorized by relative level of chronic local human disturbance (detailed in Table S1), and villages (pink circles) are scaled to human population size. Sites at which high-precision *in situ* temperature loggers were deployed between 2011 and 2017 are circled in black. Inset shows Kiritimati's location in the central equatorial Pacific Ocean.

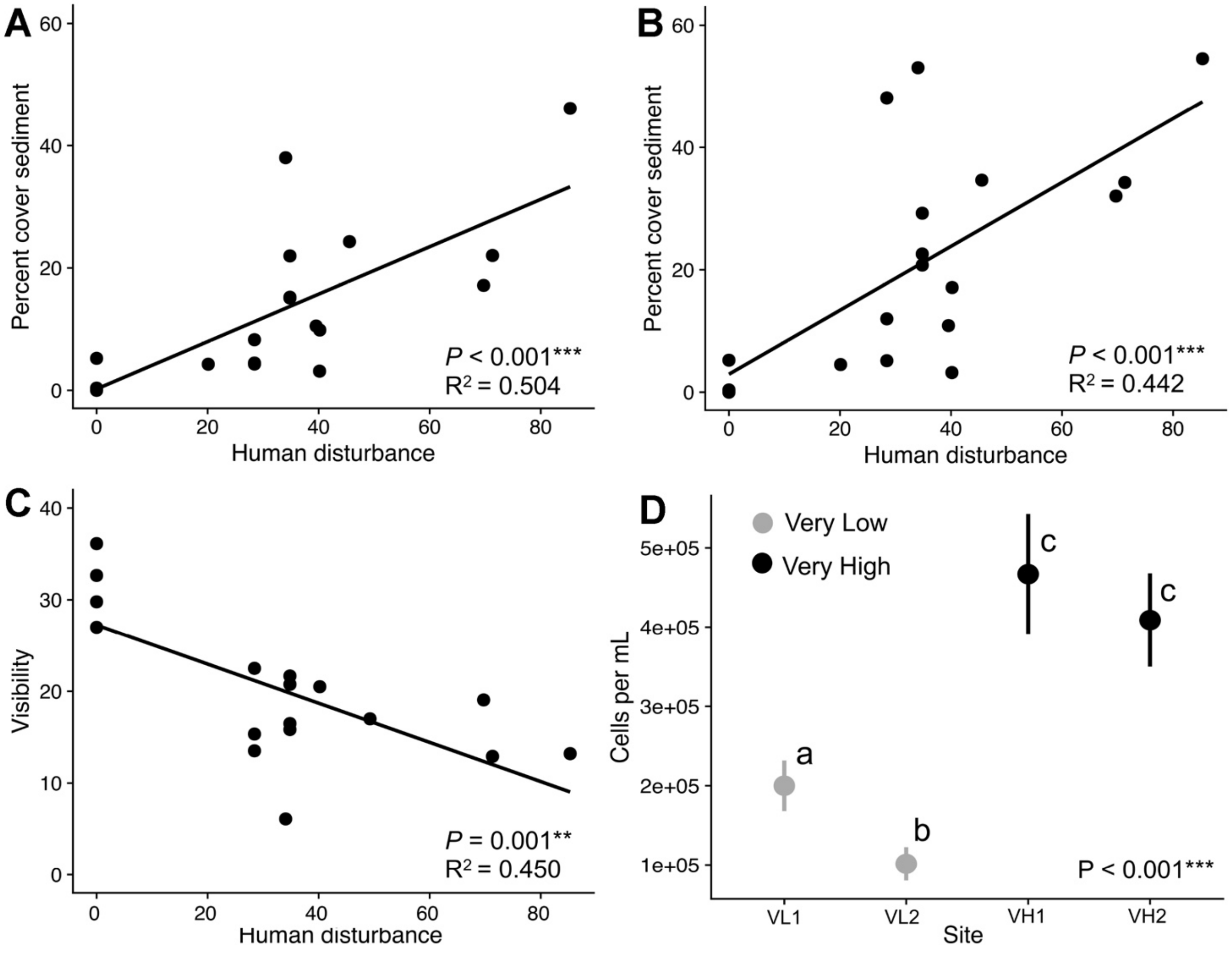


**Fig. S3. Indicators of human disturbance across Kiritimati atoll.** **A** and **B**) Relationship between benthic sediment cover, both without (**A**) and with (**B**) sand included, and chronic local human disturbance (sqrt(LocalDisturb)); (**C**) Relationship between water column visibility (a proxy for turbidity) and the human disturbance index; (**D**) Comparison of microbial counts at two very high disturbance and two very low disturbance sites; data from McDevitt-Irwin *et al.* (*48*). Letters indicate significant differences between means as determined by a Tukey post-hoc test.

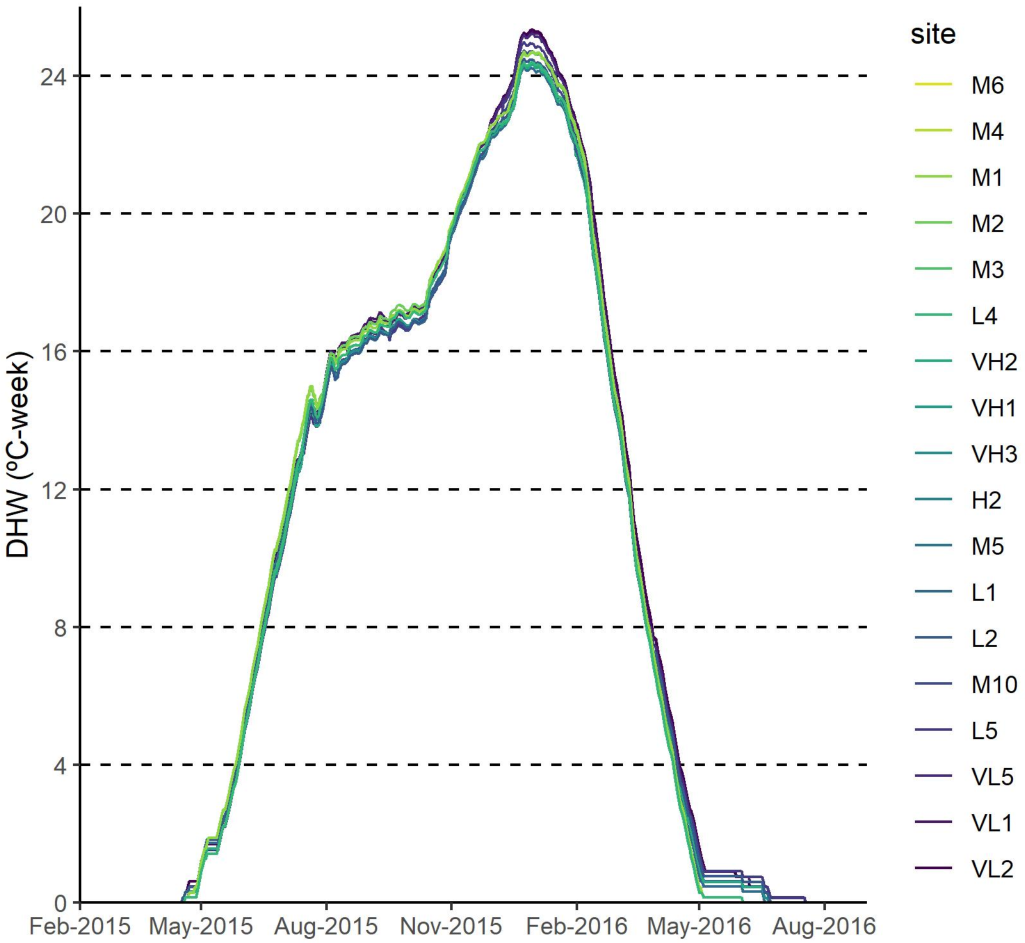


**Fig. S4. Thermal stress at each site on Kiritimati atoll during the 2015–2016 El Niño event**, measured in degree heating weeks (DHW, °C-weeks) from NOAA Coral Reef Watch (CRW). Site colors are ordered clockwise from the southwest side of the island.

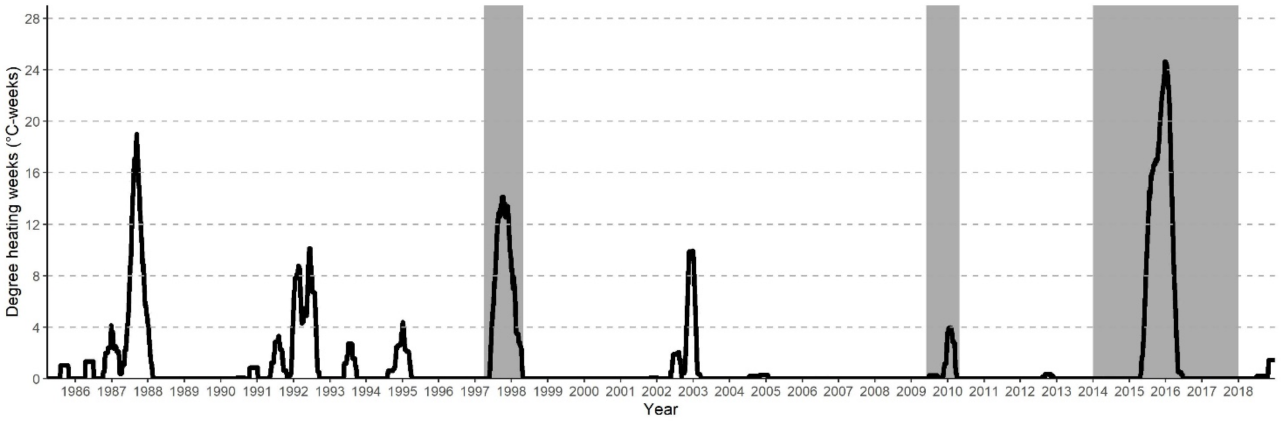


**Fig. S5. Long-term thermal stress on Kiritimati**, plotted as mean degree heating weeks (DHW, °C-weeks; across all nineteen study sites) experienced over the last thirty-four years (1985–2019) from NOAA Coral Reef Watch's (CRW) satellite-derived data product. Grey shaded areas denote the timing of the first global coral bleaching event (caused by the 1997–1998 El Niño), the second global coral bleaching event (caused by the 2009–2010 El Niño), and the third (2014–2017) global coral bleaching event (*11*).

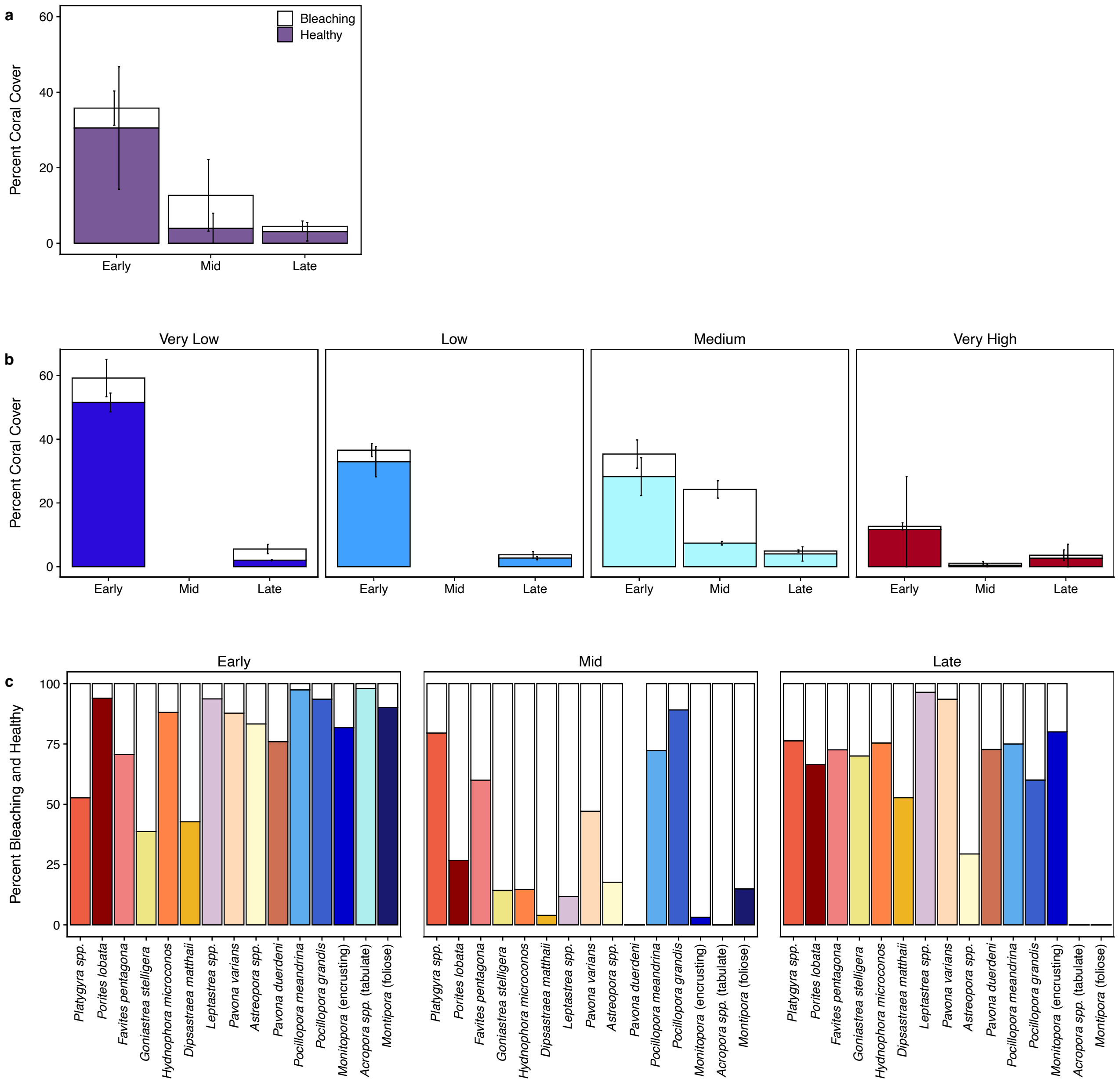


**Fig. S6. Extent of bleached coral at three time points during the prolonged 2015–2016 heatwave.** Early = July 2015 (two months heat stress); Mid = November 2015 (six months heat stress); Late = late March/April 2016 (ten months heat stress). Percent bleaching vs. healthy hard coral cover (out of total benthic community composition) (**A**) across all sites, (**B**) across sites within each disturbance category, and (**C**) for the fifteen most common hard coral species on Kiritimati (prior to the El Niño), averaged across the atoll, showing the progression of bleaching across the three time points. Species are ordered left to right from highest to lowest overall survival on the atoll by end of the heatwave. Plots include the 14 sites that were sampled during the heatwave (very low (VL1–VL3), low (L1, L2, L4), medium (M1–M5), high (none), very high (VH1–VH3)); the remaining five sites were only sampled before and after the event.

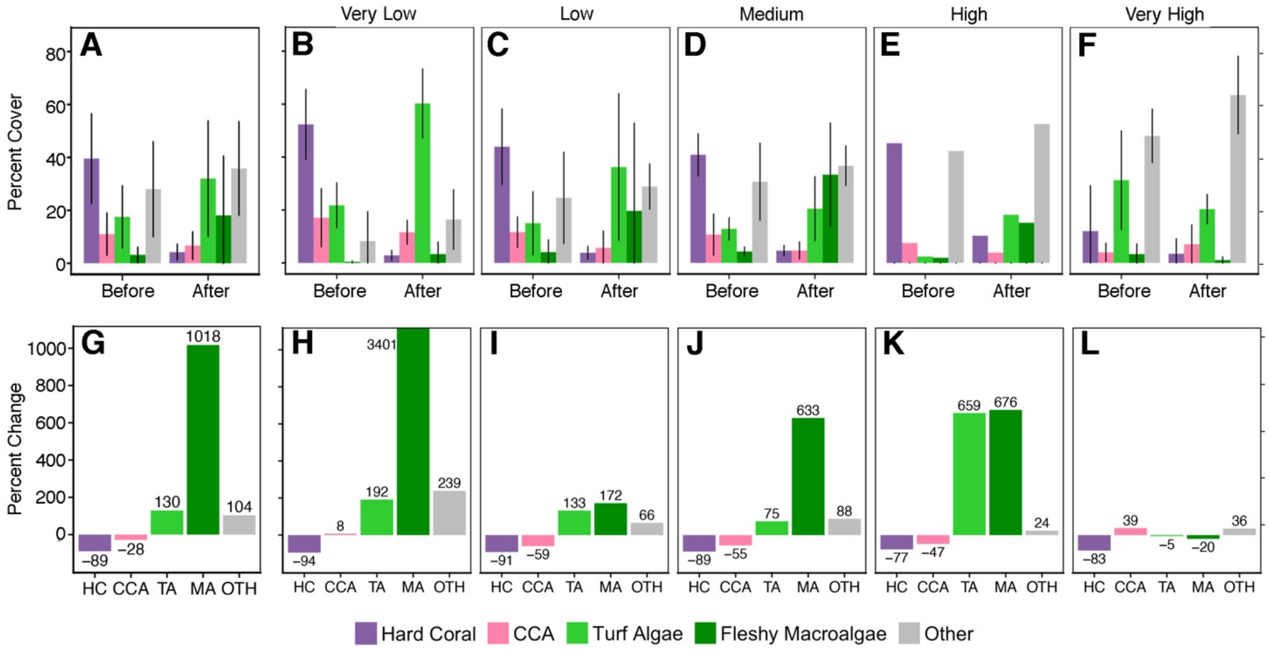


**Fig. S7. Change in overall benthic community composition on Kiritimati's reefs as a result of the prolonged 2015–2016 El Niño heatwave,** for (**A**) and (**G**) means of all sites, and (**B**–**F**, **H**–**L**) means of the sites within each of the five disturbance levels. (**A**–**F**) show percent cover of hard coral (HC), crustose coralline algae (CCA, also includes *Peyssonnelia* spp.), turf algae (TA), fleshy macroalgae (MA), and other substrates (sand, sediment, rubble, consolidated rock (i.e., exposed white calcium carbonate), soft coral) before (July 2013–May 2015) and after (November 2016, July 2017) the heatwave; (**G**–**L**) show the percent change for each benthic community component over this period.

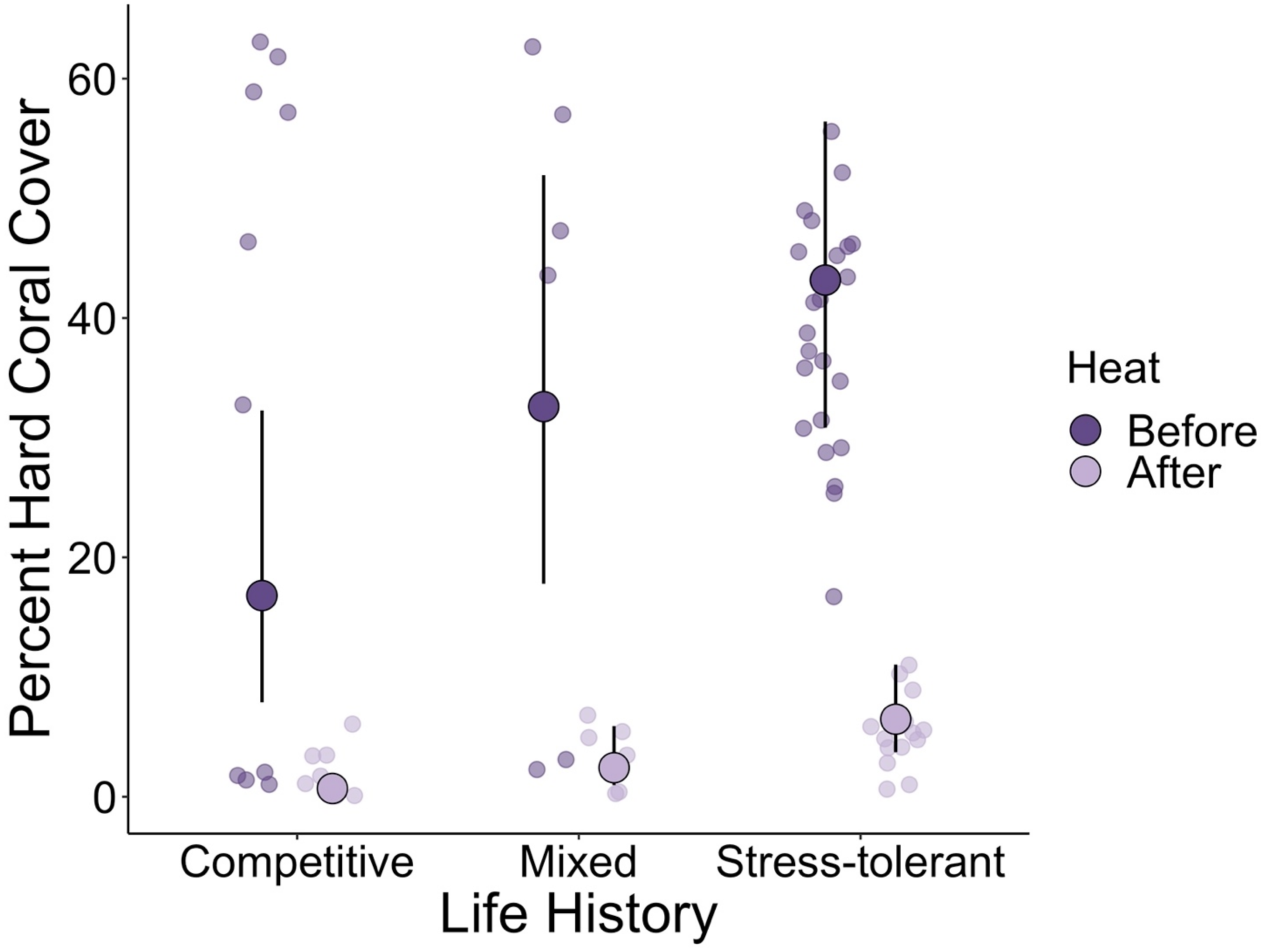


**Fig. S8. Estimated coral cover at sites dominated by competitive or stress-tolerant coral species (or 'mixed' sites with no dominant life history type) before and after the 2015–2016 El Niño.** Larger points represent predicted values (mean ± 95% confidence interval) extracted from the model, while smaller points represent observed values.

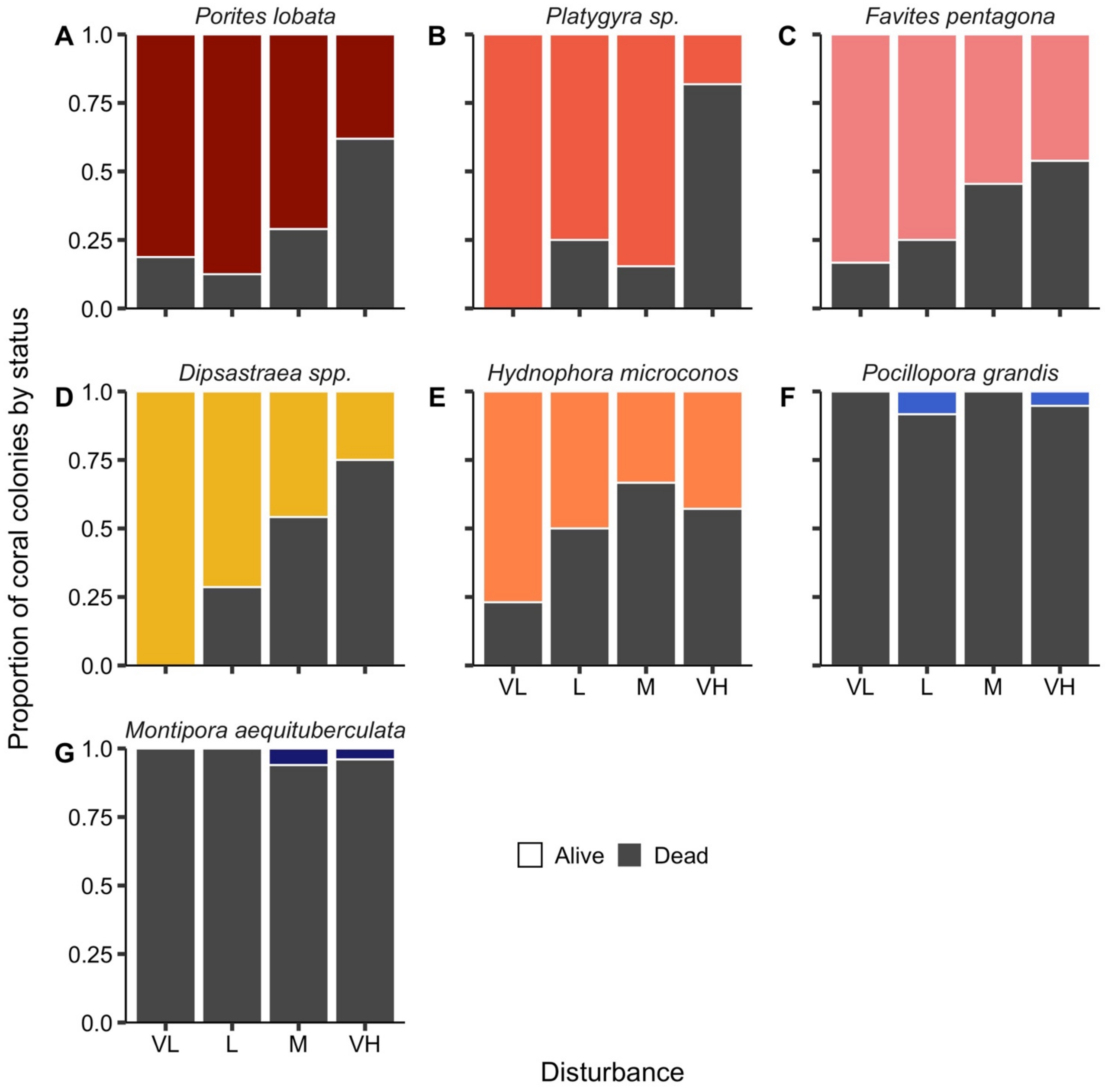


**Fig. S9. Proportion of coral colonies of each species that survived** (with species colour-coded as in Fig. 5f and Fig. S10 or died (grey), with sites categorized by chronic local disturbance (VL = very low; L = low; M = medium; VH = very high).

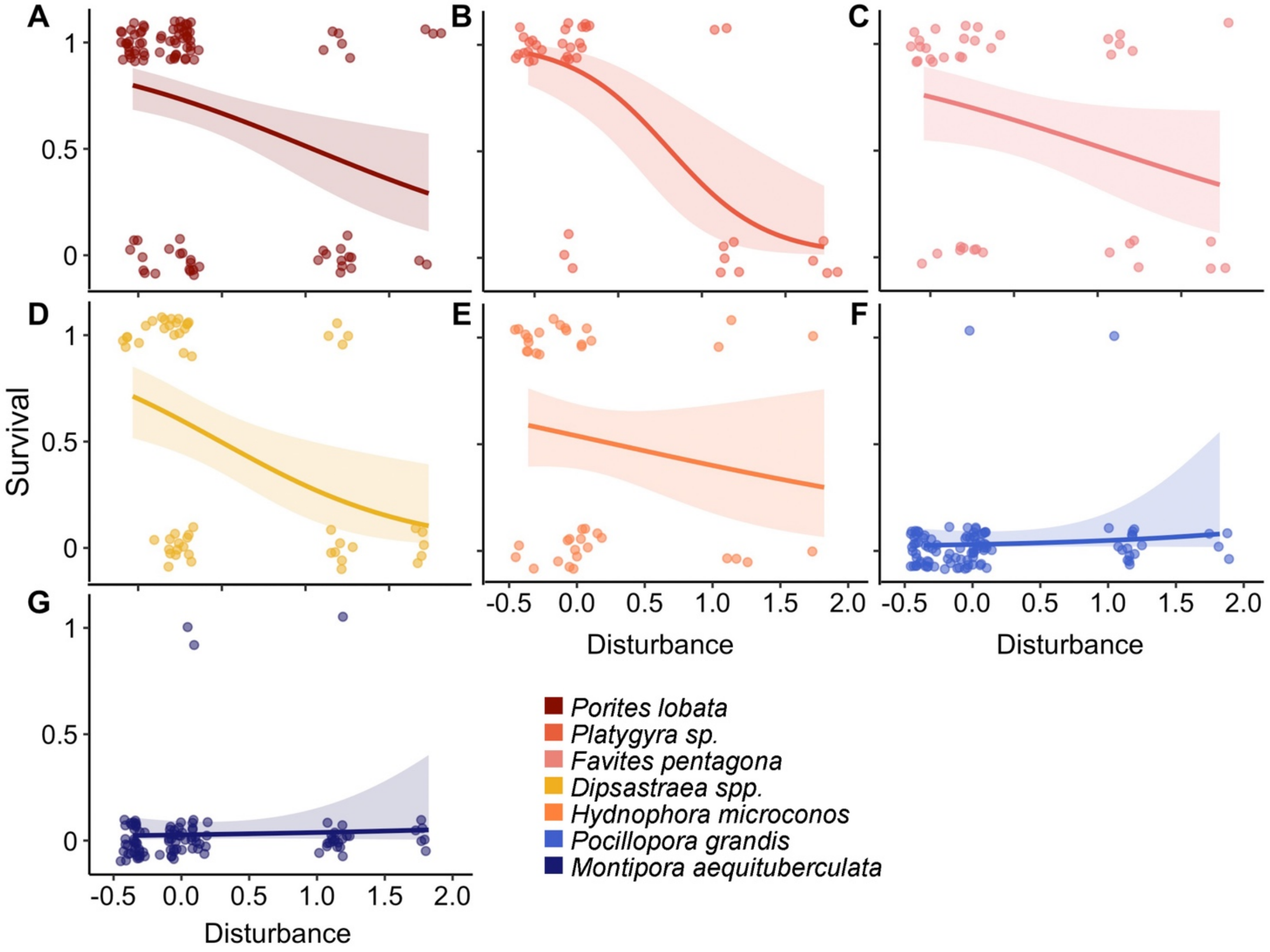


**Fig. S10. Relationship between survival (= 1 vs. 0 = died) of individual coral colonies and chronic local disturbance, by species**. Circles are individual colonies (points were x and y jittered for visualization), solid lines are the logistic regression estimate with shading denoting the 95% confidence intervals.

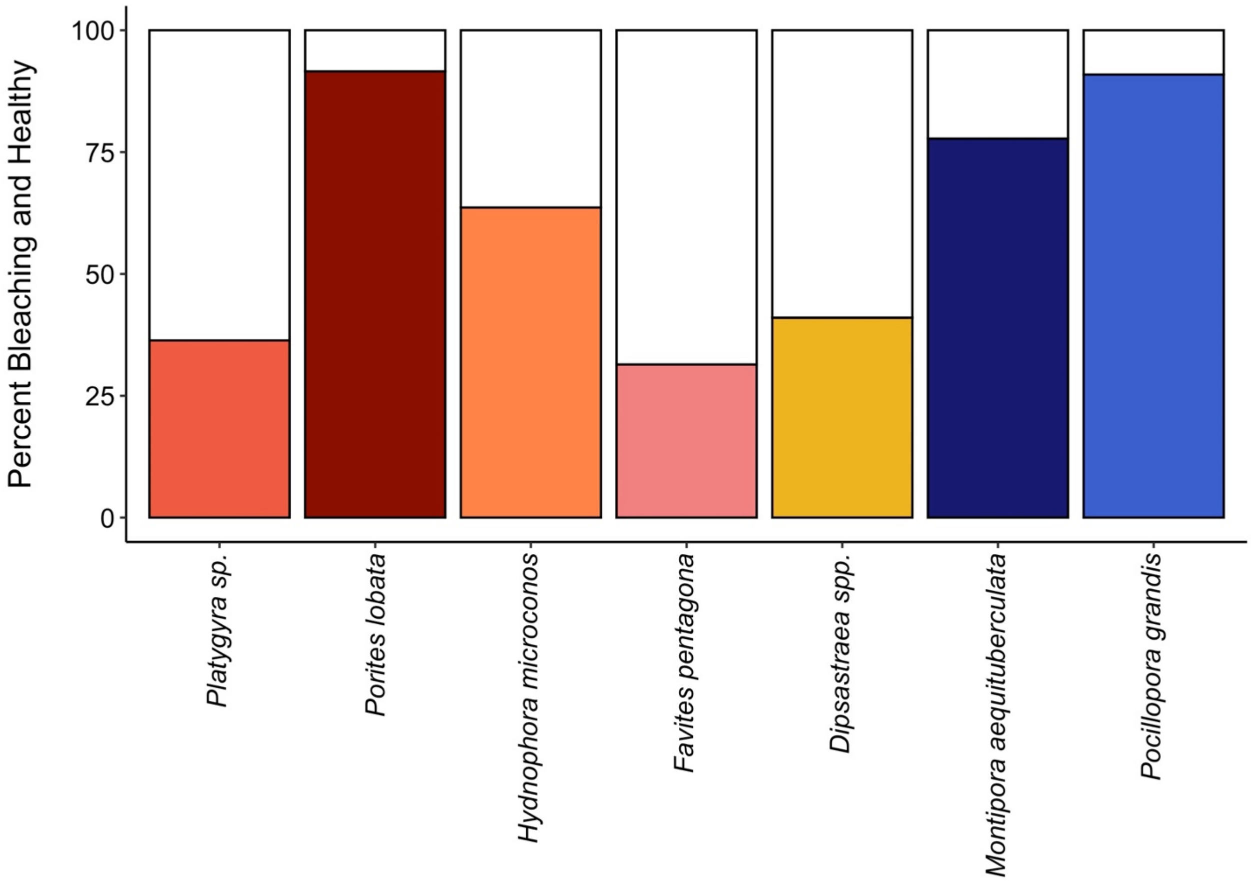


**Fig. S11. Incidence of bleaching in tagged coral colonies early in the El Niño (July 2015).** Percent bleaching (white portion) vs. healthy (colored portion) in tagged coral colonies across all sites on Kiritimati. Species are ordered left to right from highest to lowest overall survival (using tagged coral data) on the atoll by end of the heatwave.

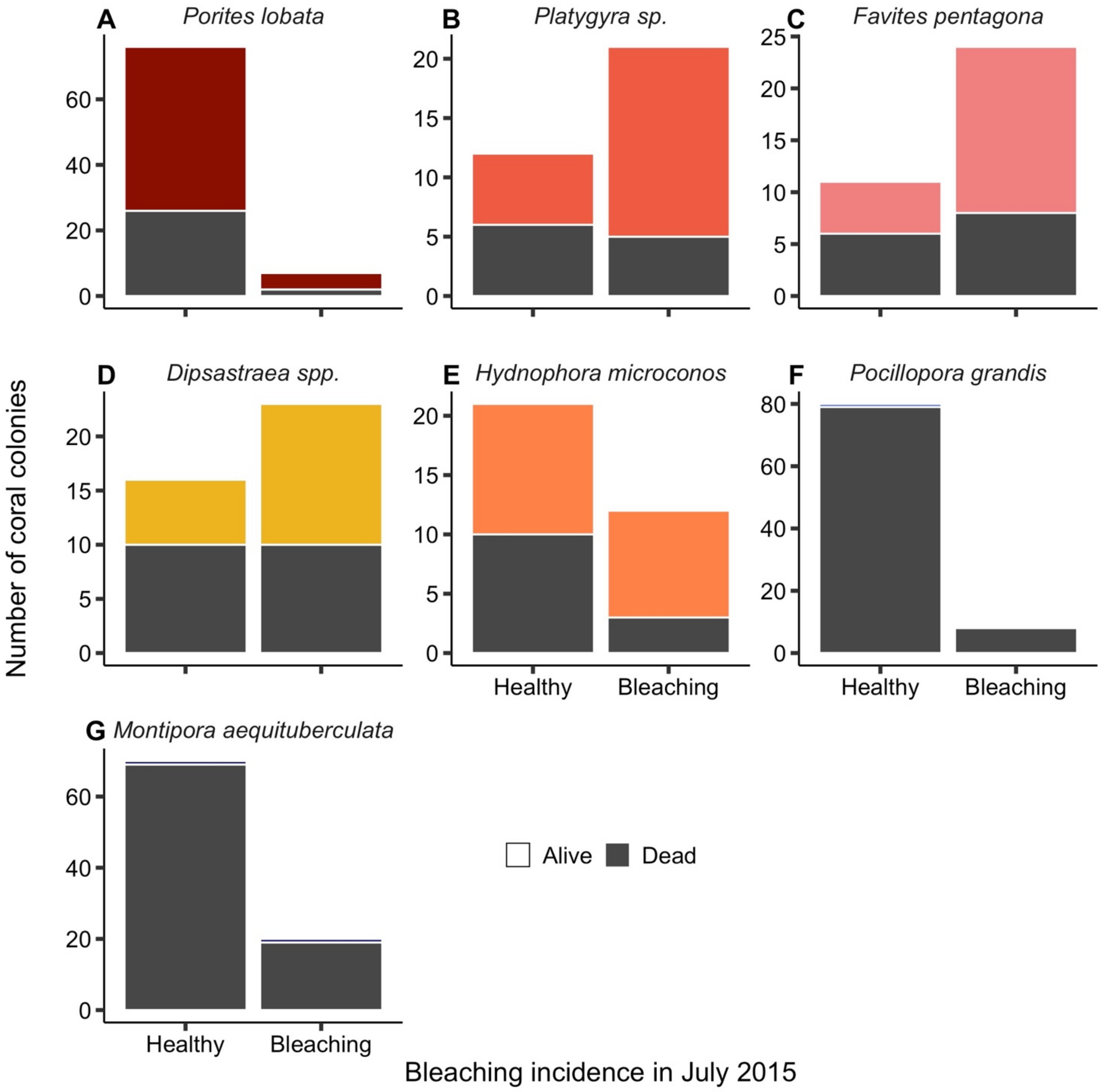


**Fig. S12. Survival status of tagged coral colonies that were either bleached or healthy early in the El Niño (July 2015).** Dead = grey; Alive = colored.

**Table S1. Chronic local disturbance at each of nineteen monitoring sites on Kiritimati.** Population is the number of people residing within 2 km of the site. Fishing pressure is the extracted value from a kernel density function of fishing pressure (*82*). Combined metric is the sum of population and fishing pressure, and sites are ordered from greatest to least disturbance according to this metric. Site numbers and disturbance level colours match those on Figure S1.

| Site | Population | Fishing Pressure | Combined Metric | Disturbance Category |
|---|---|---|---|---|
| VH1 | 4042 | 3234 | 7276 | Very High |
| VH3 | 3065 | 2021 | 5086 | Very High |
| VH2 | 1223 | 3638 | 4861 | Very High |
| H2 | 458 | 1617 | 2075 | High |
| M5 | 0 | 1617 | 1617 | Medium |
| M10 | 1209 | 404 | 1613 | Medium |
| M6 | 351 | 1213 | 1564 | Medium |
| M1 | 0 | 1213 | 1213 | Medium |
| M2 | 0 | 1213 | 1213 | Medium |
| M3 | 0 | 1213 | 1213 | Medium |
| M4 | 351 | 809 | 1160 | Medium |
| L1 | 0 | 809 | 809 | Low |
| L2 | 0 | 809 | 809 | Low |
| L4 | 0 | 809 | 809 | Low |
| L5 | 0 | 404 | 404 | Low |
| VL1 | 0 | 11 | 11 | Very Low |
| VL2 | 0 | 11 | 11 | Very Low |
| VL3 | 0 | 11 | 11 | Very Low |
| VL5 | 0 | 11 | 11 | Very Low |

**Table S2. Oceanographic characteristics of monitoring sites on Kiritimati**: Max NPP = maximum net primary productivity (mg C $m^{-2}$ $day^{-1}$), MMM = maximum monthly mean sea surface temperature (°C), Expos. = Exposure (W = windward; S = sheltered), Wave = mean wave energy (kW $m^{-1}$). Salinity (ppt), DO = dissolved oxygen (mg $L^{-1}$), pH, and phosphate (µM) are shown as means (± SE) averaged across measurements taken *in situ* in each expedition. Data sources and collection methods detailed in text. Sites are numbered and colour coded by disturbance level (as in Table S1 and Fig. S1).

| Site | Max NPP (mg C/($m^2$ day) | MMM (°C) | Expos. | Wave (kW/m) | Salinity (ppt) | DO (mg/L) | pH | Phosphate (µM) |
|---|---|---|---|---|---|---|---|---|
| VH1 | 1112.00 | 28.02 | S | NA | 35.48 ± 0.227 | 5.788 ± 0.131 | 7.999 ± 0.015 | 0.414 ± 0.026 |
| VH3 | 1097.18 | 28.02 | S | 24.95 | 35.73 ± 0.231 | 5.595 ± 0.106 | 7.974 ± 0.049 | 0.440 ± 0.057 |
| VH2 | 1158.56 | 28.02 | S | NA | 35.65 ± 0.334 | 5.659 ± 0.055 | 7.928 ± 0.022 | 0.545 ± 0.050 |
| H2 | 1097.18 | 28.02 | S | 24.95 | 35.15 ± 0.175 | 5.282 ± 0.182 | 7.988 ± 0.012 | 0.276 |
| M5 | 1035.06 | 28.03 | W | 25.36 | 35.37 ± 0.043 | 5.640 | 7.997 ± 0.013 | 0.406 ± 0.012 |
| M10 | 979.54 | 28.01 | W | 26.23 | 35.32 | 5.677 | 7.993 | NA |
| M6 | 984.27 | 27.99 | S | 24.80 | 35.34 ± 0.175 | 5.977 ± 0.258 | 7.891 ± 0.091 | 0.412 ± 0.412 |
| M1 | 1077.88 | 27.99 | S | 24.82 | 35.12 ± 0.101 | 5.973 ± 0.113 | 7.785 ± 0.130 | 0.446 ± 0.024 |
| M2 | 1077.88 | 27.99 | S | 24.82 | 35.28 ± 0.089 | 5.907 ± 0.124 | 7.977 ± 0.017 | 0.492 ± 0.026 |
| M3 | 1070.26 | 28.01 | S | NA | 36.41 ± 0.331 | 5.842 ± 0.130 | 7.937 ± 0.015 | 0.409 ± 0.031 |
| M4 | 984.27 | 27.99 | S | 24.80 | 36.23 ± 0.772 | 5.763 ± 0.180 | 7.954 ± 0.026 | 0.433 ± 0.038 |
| L1 | 1035.06 | 28.03 | W | 25.36 | 35.18 ± 0.170 | 6.192 ± 0.180 | 8.052 ± 0.026 | 0.417 ± 0.038 |
| L2 | 992.54 | 28.02 | W | 25.73 | 35.29 ± 0.258 | 5.397 ± 0.218 | 7.936 ± 0.029 | 0.382 ± 0.382 |

| | | | | | | | | |
|---|---|---|---|---|---|---|---|---|
| L4 | 1126.02 | 28.01 | S | NA | 35.35 ± 0.300 | 5.867 ± 0.242 | 7.952 ± 0.040 | 0.448 ± 0084 |
| L5 | 979.54 | 28.00 | W | 26.23 | 35.27 | 5.933 | 7.983 | NA |
| VL1 | 1003.66 | 27.96 | W | 26.41 | 35.14 ± 0.158 | 5.718 ± 0.128 | 7.970 ± 0.029 | 0.448 ± 0.038 |
| VL2 | 993.43 | 27.96 | W | NA | 35.06 ± 0.148 | 5.737 ± 0.383 | 8.013 ± 0.032 | 0.468 ± 0.052 |
| VL3 | 1031.55 | 27.97 | S | 24.90 | 35.58 ± 0.693 | 5.581 ± 0.061 | 7.938 ± 0.013 | 0.443 ± 0.036 |
| VL5 | 1015.12 | 27.97 | W | 26.07 | 35.11 | 7.000 | 7.977 | 0.453 |

**Table S3. Comparison of maximum thermal stress** (degree heating weeks) and maximum temperature anomaly (above the mean monthly maximum temperature, MMM) across sites on Kiritimati (from NOAA's CRW product (*11*)) during the 2015–2016 El Niño event.

| Site | Maximum Thermal Stress (degree heating weeks, °C-weeks) | Maximum Temperature Anomaly (°C above MMM) |
|---|---|---|
| VH1 | 24.36 | 2.86 |
| VH3 | 24.36 | 2.86 |
| VH2 | 24.33 | 2.87 |
| H2 | 24.36 | 2.86 |
| M5 | 24.26 | 2.83 |
| M10 | 24.75 | 2.92 |
| M6 | 24.71 | 2.89 |
| M1 | 24.71 | 2.89 |
| M2 | 24.69 | 2.90 |
| M3 | 24.39 | 2.87 |
| M4 | 24.71 | 2.89 |
| L1 | 24.26 | 2.83 |
| L2 | 24.5 | 2.88 |
| L4 | 24.39 | 2.87 |
| L5 | 24.99 | 2.94 |
| VL1 | 25.28 | 3.04 |
| VL2 | 25.34 | 3.05 |
| VL3 | 25.05 | 2.95 |
| VL5 | 25.2 | 3.02 |
| **Mean** | **24.67** | **2.91** |
| **Std. Dev.** | **0.36** | **0.07** |

**Table S4**. **Results (parameter estimates) for fixed effects from generalized linear mixed-effects models examining the influence of benthic and environmental variables on hard coral cover before the heatwave** (**A**) and across heatwave periods (**B**), and on the loss of coral cover following the heatwave (**C**). Bold indicates significant difference from baseline levels (Exposure = Windward; Heat = Before; Life History = Stress-tolerant) at α = 0.05; asterisks indicate levels of significance (* $p < 0.05$, ** $p < 0.01$, *** $p < 0.001$). Red shaded boxes denote variables with negative estimates, indicating a decline compared to baseline levels.

A)

| Model | n | Disturbance | NPP | MMM | Exposure |
|---|---|---|---|---|---|
| Model 1 | 39 | **-1.987*****| -0.379 | 0.217 | -0.064 |
| Model 1 | 19 | **-1.738***** | -0.315 | 0.152 | 0.015 |

B)

| Model | n | Heat | Disturbance | Life History | | DHW | Heat* Disturbance | Heat*Life History | |
|---|---|---|---|---|---|---|---|---|---|
| | | | | Mixed | Competitive | | | Mixed | Competitive |
| Model 2 | 67 | **-2.463***** | **-2.296***** | - - - | - - - | -0.155 | **1.567***** | - - - | - - - |
| Model 2 | 38 | **-2.684***** | **-2.101***** | - - - | - - - | -0.211 | **1.507***** | - - - | - - - |
| Model 3 | 67 | **-2.395***** | - - - | -0.452 | **-1.324*** | **1.491***** | - - - | -0.574 | **-0.975**** |
| Model 3 | 38 | **-2.488***** | - - - | -0.398 | **-1.161*** | **1.438**** | - - - | -0.545 | **-0.949*** |
| Model 4a | 67 | **-3.694***** | **-1.493***** | - - - | - - - | **0.736*** | **2.057***** | - - - | - - - |
| Model 4a | 38 | **-4.949***** | **-1.432***** | - - - | - - - | **0.798*** | **1.869***** | - - - | - - - |
| Model 4b | 67 | **-1.665***** | **-2.300***** | - - - | - - - | **-1.006**** | **0.908***** | - - - | - - - |
| Model 4b | 38 | **-1.803***** | **-1.866***** | - - - | - - - | **-1.046**** | **0.802*** | - - - | - - - |
| Model 5a | 67 | **-2.417***** | 0.731 | - - - | - - - | **1.286***** | **2.170***** | - - - | - - - |
| Model 5a | 38 | **-2.786***** | 0.581 | - - - | - - - | **1.370***** | **1.520**** | - - - | - - - |
| Model 5b | 67 | **2.894***** | -0.423 | - - - | - - - | **-0.680*** | **0.893*** | - - - | - - - |
| Model 5b | 38 | **2.592***** | -0.259 | - - - | - - - | **-0.713*** | 0.755 | - - - | - - - |

C)

| Model | n | Proportion bleached | Disturbance | DHW |
|---|---|---|---|---|
| Model 6 | 13 | -0.207 | 0.044 | **0.917*** |
| Model 7a | 13 | -0.092 | 1.051 | 1.496 |
| Model 7b | 13 | -0.042 | -0.219 | 0.324 |

*Note:* NPP = maximum net primary productivity (mg C $m^{-2}$ $day^{-1}$); MMM = maximum monthly mean temperature (°C); DHW = degree heating weeks (°C-weeks). Models in (**A**) and (**B**) were fit using two different data sets: one including all values such that some sites had more than one data point per heatwave period (n = 39 [**A**]; n = 67 [**B**]) and one where values were averaged to produce one data point per site for each heatwave period (n = 19 [**A**]; n = 38 [**B**]). In Model 4, the cover of each life-history type is calculated as the proportion of overall benthic cover, while in Model 5 it is calculated as the proportion of total hard coral cover.

Model structures are as follows:

**A**) Model 1: Overall coral cover ~ Disturbance + NPP + MMM + Exposure
**B**) Model 2: Overall coral cover ~ Heat * Disturbance + DHW
Model 3: Overall coral cover ~ Heat * Life history + DHW
Model 4/5a: Competitive coral cover ~ Heat * Disturbance + DHW
Model 4/5b: Stress-tolerant coral cover ~ Heat * Disturbance + DHW
**C**) Model 6: Loss of overall hard coral cover ~ Proportion bleached + Disturbance + DHW
Model 7a: Loss of competitive coral cover ~ Proportion bleached + Disturbance + DHW
Model 7b: Loss of stress-tolerant coral cover ~ Proportion bleached + Disturbance + DHW

**Table S5. Model results (parameter estimates) from logistic regression models examining influences of survival on tagged coral colonies.** Bold indicates significant difference from baseline levels (i.e., stress-tolerant, *Porites lobata*) at $\alpha = 0.05$; asterisks indicate levels of significance (* $p < 0.05$, ** $p < 0.01$, *** $p < 0.001$). Red shaded boxes denote variables with negative estimates, indicating a decline compared to baseline levels.

| | | Overall Model | Life History Model | Species Model |
|---|---|---|---|---|
| Human Disturbance Continuous | | **-0.45716 ± 0.09852*** ** | **-1.1749 ± 0.2056*** ** | **-1.0410 ± 0.3489** ** |
| Life History | Competitive | - - - | **-4.5880 ± 0.5354*** ** | - - - |
| Disturbance x Life History | | - - - | **1.7252 ± 0.6352** ** | - - - |
| Coral Species | *Platygyra ryukyuensis* | - - - | - - - | 0.8934 ± 0.6365 |
| | *F. pentagona* | - - - | - - - | -0.1948 ± 0.4549 |
| | *Dipsastraea* spp. | - - - | - - - | -0.6015 ± 0.4054 |
| | *H. microconos* | - - - | - - - | **-0.9001 ± 0.4021*** |
| | *P. grandis* | - - - | - - - | **-5.1228 ± 0.8778*** ** |
| | *M. aequituberculata* | - - - | - - - | **-4.5906 ± 0.6967*** ** |
| Disturbance x species | *Platygyra ryukyuensis* | - - - | - - - | **-1.7636 ± 0.8692*** |
| | *F. pentagona* | - - - | - - - | 0.2118 ± 0.5741 |
| | *Dipsastraea* spp. | - - - | - - - | -0.3640 ± 0.6240 |
| | *H. microconos* | - - - | - - - | 0.4780 ± 0.6565 |
| | *P. grandis* | - - - | - - - | 1.8423 ± 0.9944 |
| | *M. aequituberculata* | - - - | - - - | 1.3952 ± 0.8632 |

**Table S6. Coral taxa on Kiritimati.** Corals are categorized by the fifteen most common taxa identified in the photoquadrat images (processed using CoralNet) across the n = 19 study sides prior to the 2015–2016 El Niño, and other rarer species. The fifteen most common corals are ordered from most to least common before the El Niño and their rank and proportion after the El Niño is also given. Coral life history strategy retrieved from the Coral Traits Database release 1.1.1 (https://coraltraits.org/) (*88*), unless otherwise noted. Current taxonomy (and name synonymy) retrieved from WoRMS (http://www.marinespecies.org/).

| Life History | Family | Species | Notes | Rank and Proportion After |
|---|---|---|---|---|
| **Top 15 most common coral taxa*** | | | | |
| Stress-tolerant | Poritidae | *Porites lobata* | May include *P. evermannii* and *P. lutea* | 1 (51.4%) |
| Competitive | Acroporidae | *Montipora aequituberculata* | *M. aequituberculata* with foliose morphology | Tied 14 (0%) |
| Competitive | Acroporidae | *Montipora* spp. | *Montipora* spp. with encrusting morphology, includes *M. aequituberculata* and a few potentially unnamed species | 13 (0.3%) |
| Competitive | Pocilloporidae | *Pocillopora grandis* | Synonym: *Pocillopora eydouxi* | 6 (3.4%) |
| Stress-tolerant | Merulinidae | *Hydnophora microconos* | | 5 (3.6%) |
| Stress-tolerant | Merulinidae | *Dipsastraea matthaii* | Synonym: *Favia matthaii* | 8 (3.0%) |
| Competitive | Pocilloporidae | *Pocillopora meandrina* | | 10 (1.4%) |
| Stress-tolerant | Merulinidae | *Goniastrea stelligera* | Synonym: *Favia stelligera* | 3 (9.9%) |
| Stress-tolerant | Agariciidae | *Pavona varians* | | 11 (1.3%) |
| Competitive | Acroporidae | *Acropora hyacinthus* | Tabulate morphology | Tied 14 (0%) |
| Stress-tolerant | Merulinidae | *Platygyra* spp. | Primarily *P. ryukyuensis*, may include *P. contorta*, *P. daedalea*, *P. sinensis* | 2 (10.6%) |
| Weedy | Faviidae (synonym: Incertae sedis) | *Leptastrea* spp. | Includes *L. pruinosa and L. bewickensis* | 9 (2.9%) |
| Stress-tolerant | Acroporidae | *Astreopora* spp. | Includes *A. cucullata, A. myriophthalma*, and *A. suggesta* | 12 (1.2%) |
| Stress-tolerant | Merulinidae | *Favites pentagona* | | 4 (4.0%) |

| | | | | |
|---|---|---|---|---|
| Stress-tolerant | Agariciidae | *Pavona duerdeni* | | 7 (3.1%) |
| **Other coral taxa[*] (i.e Rare species)** | | | | **3.9%** |
| Competitive | Acroporidae | *Acropora* spp. | Corymbose morphology (*A. rosaria*, synonym: *A. loripes*; *A. subulata*; and hybrids of these species) | |
| Competitive | Acroporidae | *Acropora* spp. | Includes digitate morphology (*A. digitifera*) and any species in the genus *Acropora* that could only be identified to genus. This was often the case with coral recruits that had not yet developed distinguishing morphological characteristics. | |
| Competitive[†] | Dendrophylliidae | *Turbinaria reniformis* | | |
| Competitive[‡] | Pocilloporidae | *Pocillopora zelli* | Includes *P. zelli* and also *Pocillopora* sp. recruits that could not be identified to species (likely includes *P. meandrina* and *P. grandis* recruits). | |
| Stress-tolerant | Agariciidae | *Gardineroseris planulata* | | |
| Stress-tolerant[§] | Agariciidae | *Leptoseris mycetoseroides* | | |
| Stress-tolerant | Fungiidae | *Fungia* spp. (also *Lithophyllon* sp., *Danafungia* spp., *Pleuractis* sp., and *Lobactis* sp.) | Includes *F. concinna* (synonym: *Lithophyllon concinna*), *F. corona* (synonym: *Danafungia scruposa*), *F. danai* (synonym: *D. horrida*), *F. granulosa* (synonym: *Pleuractis granulosa), F. scutaria* (synonym: *Lobactis scutaria*) | |
| Stress-tolerant[‖] | Fungiidae | *Herpolitha limax* | | |
| Stress-tolerant[¶] | Fungiidae | *Sandalolitha robusta* | | |

| | | | |
|---|---|---|---|
| Stress-tolerant# | Lobophylliidae | *Lobophyllia hemprichii* | |
| Stress-tolerant | Merulinidae | *Dipsastraea speciosa* | Synonym: *Favia speciosa* |
| Stress-tolerant | Merulinidae | *Astrea* spp. (synonym: *Montastraea* spp.) | May include *A. annuligera* (synonym: *Montastraea annuligera*), *A. curta* (synonym: *M. curta*) |
| Stress-tolerant | Merulinidae | *Favites halicora* | |
| Generalist | Merulinidae | *Hydnophora exesa* | |

* Coral taxa included in the top fifteen comprised between 8.32% (*Porites lobata*) and 1% (0.97%; *Acropora* corymbose) of total hard coral cover prior to the El Niño, with the remaining 'Other' coral taxa each comprising less than 1% of total hard coral cover.

† Life history strategy extracted from congeneric *Turbinaria mesenterina*

‡ Life history strategy extracted from congeneric *Pocillopora eydouxi*

§ Life history strategy extracted from family Agariciidae (i.e., *Gardineroseris* and *Pavona*)

|| Life history strategy extracted from family Fungiidae (i.e., *Fungia*)

¶ Life history strategy extracted from family Fungiidae (i.e., *Fungia*)

### Life history strategy extracted from congenerics *Lobophyllia corymbosa* and *Lobophyllia pachysepta*

**Table S7. Number of small benthic photoquadrats (PQs) photographed per disturbance level on Kiritimati in each of nine expeditions straddling the 2015–2016 El Niño**: four before (2013–2015), three during (2015–2016) and two after (2016–2017) the event. Numbers in parentheses indicate the number of different sites that the PQs were photographed at in each disturbance level on each expedition.

| **Disturbance Level** | **Before El Niño** | | | | **During El Niño** | | | **After El Niño** | | **Total PQs per Disturbance Level** |
|---|---|---|---|---|---|---|---|---|---|---|
| | Jul 2013 | Aug 2014 | Jan 2015 | May 2015 | Jul 2015 | Nov 2015 | Mar 2016 | Nov 2016 | Jul 2017 | |
| Very Low | 73 (3) | 0 | 28 (1) | 44 (2) | 87 (3) | 0 | 59 (2) | 60 (2) | 102 (4) | **453** |
| Low | 78 (3) | 63 (2) | 0 | 0 | 60 (2) | 0 | 59 (2) | 0 | 88 (3) | **348** |
| Medium | 177 (7) | 135 (5) | 60 (2) | 90 (3) | 149 (5) | 36 (2) | 102 (4) | 120 (4) | 193 (7) | **1,062** |
| High | 30 (1) | 0 | 0 | 0 | 0 | 0 | 0 | 0 | 30 (1) | **60** |
| Very High | 87 (3) | 55 (2) | 60 (2) | 90 (3) | 89 (3) | 64 (2) | 90 (3) | 90 (3) | 89 (3) | **714** |
| **Total PQs (Sites) per Expedition** | **445 (17)** | **253 (9)** | **148 (5)** | **224 (8)** | **385 (13)** | **100 (4)** | **310 (11)** | **270 (9)** | **502 (18)** | **2,637** |

**Table S8. Number of tagged individual coral colonies of each species with known survivorship status, categorized by disturbance level.** Species are ordered from highest to lowest sample size.

| | Disturbance Level | | | | |
|---|---|---|---|---|---|
| **Species** | Very Low | Low | Mediu m | Very High | **Species Totals** |
| ***Competitive Life History Strategy*** | | | | | |
| *Montipora aequituberculata* | 35 | 8 | 33 | 25 | 101 |
| *Pocillopora grandis* | 34 | 12 | 35 | 19 | 100 |
| ***Stress-Tolerant Life History Strategy*** | | | | | |
| *Porites lobata* | 32 | 8 | 38 | 21 | 99 |
| *Dipsastraea* spp. *(D. matthaii)* | 6 | 7 | 24 | 16 | 53 |
| *Hydnophora microconos* | 13 | 6 | 15 | 7 | 41 |
| *Platygyra ryukyuensis* | 12 | 4 | 13 | 11 | 40 |
| *Favites pentagona* | 12 | 4 | 11 | 13 | 40 |
| **Totals** | **144** | **49** | **169** | **112** | **474** |
